# Single-cell transcriptomics reveal distinct subsets of activated dendritic cells in the tumor microenvironment

**DOI:** 10.1101/2022.09.13.507746

**Authors:** Tsun Ki Jerrick To, Samir Devalaraja, Ian W. Folkert, Li Zhai, Graham P. Lobel, Malay Haldar

**Affiliations:** Department of Pathology and Laboratory Medicine, Perelman School of Medicine, University of Pennsylvania, Philadelphia, PA, 19014, USA; Department of Surgery, Perelman School of Medicine, University of Pennsylvania, Philadelphia, PA, 19104, USA; Abramson Family Cancer Research Institute, Perelman School of Medicine, University of Pennsylvania, Philadelphia, PA, 19104, USA; Institute for Immunology, Perelman School of Medicine, University of Pennsylvania, Philadelphia, PA, 19104, USA

## Abstract

Dendritic cells (DCs) are rare in tumors where their heterogeneity remains unclear. To overcome the limitations of surface marker-based analyses, we utilized DC-reporter mice (*Zbtb46^GFP/+^*) to isolate tissue and tumor DCs and performed single-cell RNA sequencing (scRNA-seq). The known DC subsets were conserved across tissues, albeit at different frequencies. Activated and mature DCs formed a distinct cluster in both healthy and tumor tissues and displayed the hallmark DC migratory program (migratory DCs). We also identified a distinct subset of activated DCs in tumors that did not induce the migratory program, instead displaying signatures of interferon exposure (interferon-DCs or IFN-DCs). IFN-DCs were proficient in antigen-presentation, supported T cell proliferation, and expressed high levels of T cell-attracting chemokines. IFN-DCs further comprised of IFN1- and IFN2-DCs that were generated from CD11b+ DCs in response to type I or type II interferons respectively. We also identified IFN-DCs in scRNA-seq of human tumor-infiltrating leukocytes. These findings illustrate DC heterogeneity in tumors and suggest the existence of an interferon regulated ‘division of labor’ among activated DCs whereby the migratory DCs drive T cell priming in draining lymph nodes while the sessile IFN-DCs help recruit T cells and regulate their function in the tumor microenvironment.

**One Sentence Summary:** Tumors harbor an interferon-regulated activated sessile subset of dendritic cells.

## INTRODUCTION

Dendritic cells (DCs) are critical for initiating and regulating adaptive immune responses (*1–3*). Major DC subsets include plasmacytoid DCs (pDCs), classical DCs (cDCs), and monocyte-derived DCs (moDCs). While pDCs excel at producing type I interferons (IFN-I) in response to viral infections, cDCs and moDCs specialize in antigen presentation (*1*). cDCs originate from precursors in the bone marrow (BM) and comprise of two major subsets: cDC1s and cDC2s (*2*). cDC1s preferentially interact with CD8^+^ T cells and can cross-present cell-associated antigens, while cDC2s generally communicate with CD4^+^ T cells to regulate T helper cell polarization (*4*). Circulating monocytes can generate DCs during inflammation but their functions are relatively less clear (*5, 6*). Recent work have begun to reveal the complex network of DC subsets in tissue but how DC functions are distributed across the different DC subsets remains to be fully understood (*7*).

Monocytes, macrophages, and DCs comprise the mononuclear phagocyte (MNP) system (*2*). While monocytes and macrophages are abundant in most solid tumors, DCs are rare. cDC1s are particularly rare, accounting for just <0.5% of tumor-infiltrating leukocytes (TILs) in some tumors (*6, 8*). Nonetheless, cDC1 deficiency leads to loss of anti-tumor CD8 T cell responses, highlighting the ‘outsized’ impact of DC subtypes on tumor immune responses (*9*). The current consensus on DC subsets is based on flow cytometry, which is limited by the repertoire of cell surface markers. In contrast, single cell RNA-sequencing (scRNA-seq)-based analyses take into account the entire transcriptome of individual cells, making it unbiased, comprehensive, and hence better suited to uncover true cellular heterogeneity (*10*). However, most published scRNA-seq datasets of bulk tissue or tumor contain very few DCs, thus limiting our ability to identify rare DC subsets (*11*). Isolation of DCs for focused scRNA-seq is constrained by overlapping expression of cell-surface markers by related cell types, particularly macrophages, and potential alterations in epitopes during enzymatic tumor digestion for single cell suspension. Lineage-specific reporters can overcome many of these limitations, as we describe in this manuscript.

The prevailing view of tumor immunity is that intratumoral DCs mature and migrate (mature migratory DCs or mDCs) to draining lymph nodes (LN) where mDCs initiate an anti-tumor T cell response. These tumor-reactive T cells must then migrate to the tumor where their activity is further influenced by the suppressive or stimulatory effects of intratumoral MNPs. In this context, the suppressive effects of tumor-associated macrophages (TAMs) and myeloid-derived suppressor cells (MDSCs) are well established. However, how DCs might modulate T cell function in an antigen-specific manner within the tumor microenvironment (TME) is less clear. In this study, we utilized the *Zbtb46*^GFP/+^ reporter mice to isolate and perform scRNA-seq on DCs from normal lungs and tumors (sarcomas and melanoma). Our analyses confirm several key findings from recent studies of DC heterogeneity and identifies a distinct activated DC subset (IFN-DCs) in tumors. We also provide insights into the origin, function, and regulation of IFN-DCs in the TME.

## RESULTS

### Single cell RNA sequencing using reporter mice resolves DC heterogeneity in tumor

MNPs are developmentally, functionally, and phenotypically related (*12*). Lineage tracing and genetic reporter mice for accurate identification of specific immune cells have been developed, for example with *Lyz2^Cre^*:*Rosa26*^LSL-tdT^ to mark cells of monocyte lineage or *Zbtb46*^GFP/+^ to identify *bona fide* DCs (*13, 14*). Within inflammatory environments such as the TME, MNPs show overlap in cell surface markers, including the prototypical DC surface markers CD11c and MHCII (*15*). Correspondingly, we found that while CD11c^+^MHCII^+^ cells were dominantly Zbtb46-GFP^+^ in lymphoid tissues such as the spleen, CD11c^+^MHCII^+^ cells were heterogeneous for Zbtb46-GFP expression in murine fibrosarcomas (FS tumors; generated by syngeneic transplantation of a methylcholanthrene-induced fibrosarcoma cell line) (Figure 1A). The Zbtb46-GFP^+^ subset of CD11c^+^MHCII^+^ cells in the FS tumor was largely negative for the macrophage marker F4/80 and divided into distinct CD11b^+^ and CD103^+^ populations characteristic of peripheral tissue DCs (Figure 1A). Conversely, the Zbtb46-GFP^-^ subset of CD11c^+^MHCII^+^ cells contained a sizeable F4/80^+^ population (Figure 1A). At a functional level, the different subsets of intratumoral CD11c^+^MHCII^+^ cells based on Zbtb46-GFP and F4/80 expression varied in their ability to promote proliferation of αCD3/CD28-activated T cells (Figure 1B). While Zbtb46-GFP^+^ F4/80^-^ cells significantly promoted the proliferation of T cells, Zbtb46-GFP^-^ cells, irrespective of F4/80 expression, did not. Hence, Zbtb46-GFP expression served as a useful reporter for stimulatory DCs in the tumor (tuDCs).

**Figure 1:**
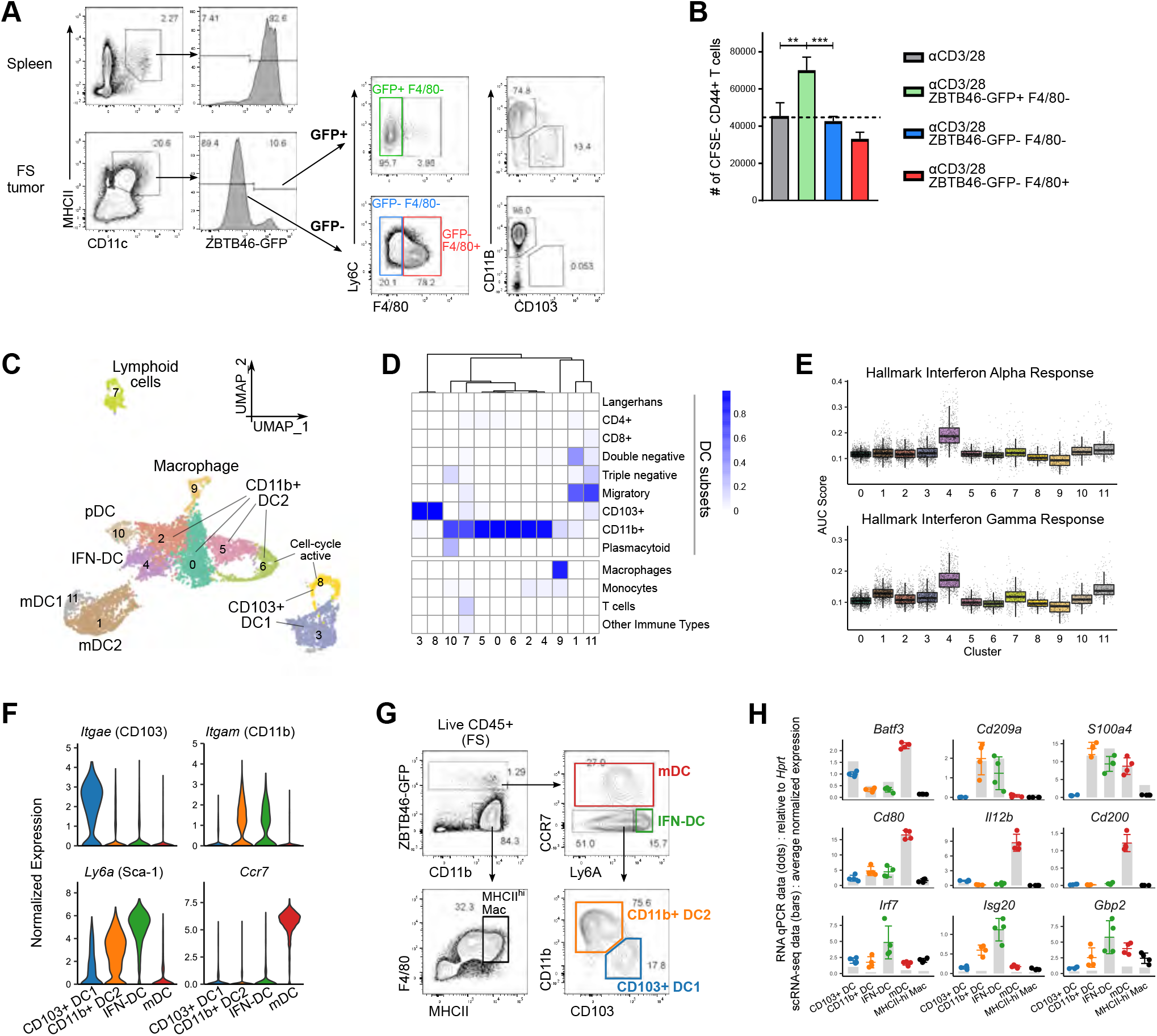
DC-focused scRNA-seq using reporter mice reveals intratumoral DC heterogeneity. A) Flow cytometry profiling of the distribution of antigen-presenting cells (APCs) within MHCII+CD11c+ cells in the spleen and a FS tumor in ZBTB46-GFP hosts. Events were pre-gated on live CD45+ cells. B) Proliferation of αCD3/CD28-activated CFSE-labeled T cells co-cultured for 3 days with the indicated tumor APCs. Tumor APCs were sorted from FS tumors in ZBTB46-GFP hosts based on the gating strategy from (A). C) UMAP plot of merged scRNA-seq (10x genomics) data of 7,490 sorted live CD45+ZBTB46-GFP+*Lyz2*-tdT+ and *Lyz2*-tdT-cells from pooled FS tumors (see Fig S1A). Shown are the numbered clusters from Seurat analyses as well as cell identities based on expression of marker genes and gene sets. D) Heatmap displaying annotation of cells in the FS tuDC scRNA-seq dataset using reference gene expression data from ImmGen. Scores represent the proportion of cells within a cluster assigned to a specific cell-type using the *SingleR* analyses tool. E) Boxplots comparing the relative enrichment of interferon response hallmark gene sets among the clusters in FS tuDC scRNA-seq dataset as calculated using *AUCell* analyses tool. F) Violin plots showing the relative expression of marker genes across the four major tuDC in the FS tuDC scRNA-seq dataset: CD103+ DC1 (c3 and c8), CD11b+ DC2 (c0, c2, c5, c6), IFN-DC (c4) and mDC (c1, c11). G) Gating strategy to identify the major tuDC subests from the scRNA-seq dataset. Events are pre-gated on live CD45+ cells from FS tumor in ZBTB46-GFP hosts. H) Expression of the indicated genes (header) within each cell population (Y-axis) in scRNA-seq dataset (grey bars) and RT-qPCR (colored dots; each dot represents one biological replicate) performed on the corresponding cells isolated by the gating strategy in (G). For each gene, the averaged normalized single-cell expression values were scaled with respect to the highest mean qPCR value. \*\**p* < 0.01, \*\*\**p* < 0.001. Statistical analysis was performed with unpaired *t* test (B). Data are shown as mean ± SD.

cDC1s are easily identified by flow cytometry (FC) based on the expression of characteristic surface markers including CD103, XCR1, and CLEC9a. In contrast, non-cDC1s show significant phenotypic and ontological heterogeneity, comprising of additional subsets of cDC2s and moDCs. Lineage tracing with *Lyz2*^Cre^ (lysozyme 2^Cre^; *LysM*^Cre^) has been widely used to examine monocyte-derived cells including DCs (*16*). However, *Lyz2* is widely expressed in myeloid cells and the utility of *Lyz2*^Cre^ in distinguishing cDC2s from moDCs in tumors is unclear. We thus generated *Zbtb46*^GFP/+^:*Lyz2*^Cre^:*Rosa26*^LSL-tdT^ mice that expressed tdTomato (tdT) in *Lyz2*-expressing cells (and their progenies) along with GFP in DCs. Here, GFP^+^ tuDCs separated into distinct tdT^-^ and tdT^+^ fractions (Figure S1A). To examine further these two fractions and the overall population of Zbtb46-GFP^+^ tuDCs, we performed scRNA-seq (10X Genomics) of Zbtb46-GFP^+^tdT^-^ and Zbtb46-GFP^+^tdT^+^ cells from FS tumors. The two samples were pooled into one dataset and analyzed with the Seurat pipeline (*17*). We observed comparable distribution of tdT^-^ and tdT^+^ cells within the UMAP space and among unsupervised clusters (Figures S1B and S1C). Hence, contrary to our expectation, the *Lyz2*-lineage did not represent distinct tuDC subsets.

We then analyzed the overall Zbtb46-GFP^+^ tuDC population by treating both tdT^-^ and tdT^+^ fractions as one dataset. Clusters were assigned identities by comparing against gene expression profiles of DC subsets in the ImmGen database (*18*) (Figure 1C and 1D). Minor clusters displaying gene expression characteristics of lymphoid cells (cluster 7), macrophages (cluster 9) and plasmacytoid DCs (cluster 10) were excluded from further analyses. cDC1s, CD11b^+^ DC2s (encompassing both cDC2s and moDC2s), and migratory-DCs (mDCs) comprised the major tuDC clusters (Figure 1C). Subclusters within both cDC1s (cluster 8) and CD11b^+^ DC2s (cluster 6) expressed high levels of cell cycle-associated genes suggestive of proliferation (Figures 1C and S1D). CD11b^+^ DC2s (clusters 0, 2, 4, 5, 6) displayed the greatest heterogeneity and contained a distinct subcluster (cluster 4) marked by enriched type I (IFN-I) and type II (IFN-II) interferon signaling signatures (Figures 1C, 1E and S1D). Hence, we designated this interferon signaling-enriched cluster as interferon-DCs (IFN-DCs). mDCs have been previously described in scRNA-seq datasets as mregDC, *LAMP3*^+^ DC, and DC3 (*19–21*). Migration from tissue to draining nodes is a hallmark of all cDCs. Correspondingly, we found that mDCs form two further subclusters consistent with cDC1 (cluster 11, mDC1) or cDC2 origins (cluster 1, mDC2) (Figure 1C), with mDC1s enriched for *Xcr1, Cd24a* and *Il12b* while mDC2s were enriched for *S100a4* and *Sirpa* (Figure S1E). The mDC1 cluster was smaller than the mDC2 cluster, which is consistent with the lower frequency of cDC1s compared to DC2s (Figure 1C).

Hence, we found four major tuDC subsets in our scRNA-seq analyses: cDC1s, CD11b^+^ DC2s, IFN-DCs and mDCs. These subsets were characterized by high expression of *Itgae* (CD103), *Itgam* (CD11b), *Ly6a* (Sca-1), and *Ccr7* respectively (Figure 1F). A FC gating strategy was developed to identify the four major tuDC subsets based on these markers within CD45^+^Zbtb46-GFP^+^ tuDCs: CCR7^+^ **mDCs**, CCR7^-^Ly6A^hi^ **IFN-DCs**, CCR7^-^Ly6A^int/lo^CD11b^-^CD103^+^ **cDC1s** and CCR7^-^Ly6A^int/lo^CD11b^+^CD103^-^ **DC2s** (Figure 1G). We next isolated tuDC subsets using the above gating strategy and confirmed differential expression of key genes associated with each subset by RT-qPCR (*Batf3* for cDC1s; *Cd209a* and *S100a4* for CD11b^+^ DC2s; *Irf7, Isg20* and *Gbp2* for IFN-DCs; *Cd80, Il12b* and *Cd200* for mDCs) (Figure 1H). Thus, the tuDC subsets identified by our scRNA-seq can be reliably isolated for further studies.

### IFN-DCs and mDCs represent distinct subsets of activated DCs

Using a previously published gene expression signature associated with overall DC maturation (*Maturation ON* signature) (*22*), we observed that mDCs expressed the highest levels of the *Maturation ON* signature followed by IFN-DCs (Figure 2A). DCs mature under homeostatic or immunogenic conditions (*23*). As the *Maturation ON* signature consists of genes associated with both homeostatic and immunogenic maturation, we next used another published signature that refers to genes found exclusively in homeostatic maturation but not in immunogenic maturation (*Homeostatic Maturation* signature) (*22*). The *Homeostatic Maturation* signature was almost solely enriched in mDCs (Figure 2A). This implied that mDCs represented DCs that matured under both homeostatic and immunogenic conditions, while the IFN-DC state was not associated with homeostatic maturation.

**Figure 2:**
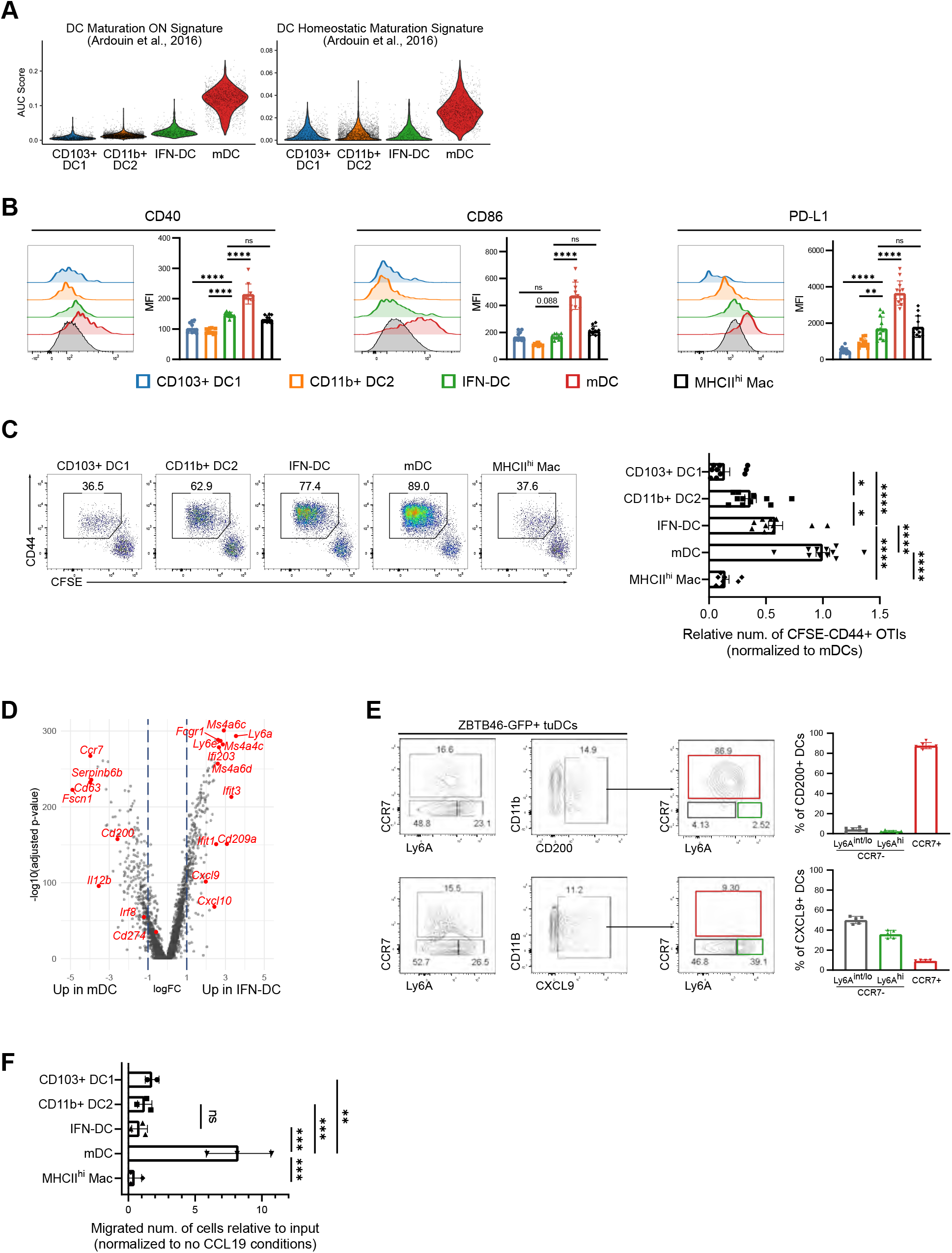
IFN-DCs and mDCs are activated DCs in the tumor microenvironment. A) Boxplots comparing the relative enrichment of the ‘DC Maturation ON’ and ‘DC Homeostatic Maturation’ signatures (from Ardouin *et al.* 2016) among the major tuDC subsets in the FS scRNA-seq dataset as calculated via *AUCell*. B) Representative histograms (left) and aggregated median fluorescent intensity (MFI, right) data for CD40, CD86 and PDL1 among FS tumor DC and macrophage subsets. C) Proliferation of OT-I T cells co-cultured with DC subsets and macrophages sorted from FS tumors and pulsed with 0.01 ng/mL SIINFEKL *ex vivo*. Representative flow cytometry gating for proliferated OT-I in each co-culture condition shown (left panel). Number of proliferated OT-I were normalized to the number of proliferated OT-I in the mDC co-culture condition and data were pooled from 3 independent experiments (right panel). D) Comparison of DEGs between mDCs (c1 and c11) and IFN-DCs (c4) from the FS scRNA-seq dataset with selected genes highlighted in red. E) FC analyses of CD200 and CXCL9 expression (bar graph) amongst various tuDC subsets (FC plots) in FS tumors Single-cell suspensions from FS tumors were incubated with brefeldin A *ex vivo* prior to intracellular staining for CXCL9. F) Migration of tuDC subsets and macrophages sorted from FS tumors across transwell membranes in response to 200 ng/mL CCL19. Number of migrated cells were normalized to no CCL19 control conditions and presented as relative to the number of input cells. Data are representative of two independent experiments. \**p* < 0.05, \*\**p* < 0.01, \*\*\**p* < 0.001, \*\*\*\**p* < 0.0001. Statistical analysis was performed with one-way ANOVA with Tukey’s post hoc test followed by multiple unpaired *t* test (B, C and F). Data are shown as mean ± SD.

DC maturation is associated with upregulation of activation (*Cd80*, *Cd86*, *Cd40*, *Relb* and *Cd83*) and immuno-modulatory genes (*Cd274*, *Pdcd1lg2*, *Cd200*, *Fas*, *Socs1*, *Socs2* and *Aldh1a2*) (*19*). Consistent with our DC maturation signature analyses above, mDCs displayed high expression across these activation and immuno-modulatory genes (Figure S2A). IFN-DCs expressed high levels of some activation markers such as *Cd40* and *Cd86*, and displayed lower expression of several immuno-modulatory genes, such as *Cd274,* compared to mDCs (Figure S2A). These findings were corroborated with FC, which showed mDCs expressing the highest levels of these surface markers (Figure 2B). IFN-DCs had higher levels of CD40 and PD-L1 compared to cDC1s and CD11b+ DC2s and trended towards higher levels of CD86 compared to CD11b^+^ DC2s (Figure 2B).

To functionally evaluate how the aforementioned maturation phenotypes translate to interactions with T cells, sorted tuDC subsets from FS tumors were pulsed with SIINFEKL and co-cultured with CFSE-labeled OT-I CD8^+^ T cells. mDCs had the greatest stimulatory capacity for inducing OT-I proliferation followed by IFN-DCs (Figure 2C). Of note, cDC1s, which were isolated based on CCR7^-^ expression, were unable to induce OT-I proliferation almost to the extent of MHCII^hi^ tumor macrophages. However, CD11b^+^ DC2s, which were also isolated based on CCR7^-^ expression, were significantly more stimulatory than cDC1s. This may imply that cDC1 activation and stimulatory capacity is tightly linked with CCR7 induction, while CD11b^+^ DC2s maintain a basal state of activation and stimulatory capacity despite CCR7^-^ expression. Finally, despite having similar levels of CD40, CD86, and PD-L1, IFN-DCs were vastly superior to MHCII^hi^ tumor macrophages in stimulating T cells. Thus, our observations at the gene and protein expression level as well as T cell co-cultures suggested mDCs were the most activated followed by IFN-DCs.

As IFN-DCs and mDCs represented activated DCs in TME, we further compared their gene expression profile to uncover potential functional specialization between the two subsets (Figure 2D). IFN-DCs were enriched for expression of CXCR3 ligands, *Cxcl9* and *Cxcl10*, that are known to recruit anti-tumor T cells to the TME (*24, 25*). Conversely, mDCs were enriched for factors such as *Il12b* and *Cd200* (Figure 2D). Consistent with this, FC of tuDCs showed CD200 expression restricted to the mDCs while CXCL9 expression was restricted to the CCR7^-^ fraction of tuDCs that contained Ly6A^hi^ IFN-DCs (Figure 2E). Additionally, genes associated with DC migration, such as *Ccr7* and *Fscn1* (*19*), were selectively expressed in mDCs but not in IFN-DCs (Figures 2D and S2B). Thus, we next examined the migratory capacity of sorted tuDC subsets from FS tumors using a transwell migration assay in response to recombinant murine CCL19, a ligand for CCR7 (Figure 2F). Consistent with their high CCR7 expression, mDCs showed the highest migratory capacity. In contrast, all other tuDC subsets, including IFN-DCs, showed limited migration. Hence, IFN-DCs are likely non-migratory activated DCs. Consistent with this notion, tumor-draining lymph nodes contained CCR7^+^ mDCs but lacked Ly6A^hi^ or CXCL9^+^ cells that could resemble IFN-DCs (Figure S2C).

To further investigate the limited migratory capacity of IFN-DCs in the transwell assay (Figure 2F), we examined the ability of the non-migratory tuDC populations to induce *Ccr7*. We found a slight induction of *Ccr7* with time in the presence of regular media in all non-migratory tuDCs (Figure S2D). However, In the presence of tumor-conditioned media, *Ccr7* was strongly upregulated in cDC1s but not in CD11b^+^ DC2s or IFN-DCs (Figure S2D). As anticipated, macrophages did not induce *Ccr7* in any condition. Thus, our results suggest that while IFN-DCs may induce *Ccr7 to some extent*, it is not sufficient to match the migratory capacity of mDCs in a transwell assay. In contrast to *Ccr7*, *Cxcl9* was downregulated in non-migratory tuDCs under all culture conditions over time (Figure S2D), which suggested that pathways regulating *Cxcl9* expression differs from that of *Ccr7*. Collectively, our results showed that IFN-DCs and mDCs were activated DC states displaying specialized functions – expression of CXCR3 ligands for IFN-DCs and migration for mDCs.

### IFN-DCs are generated in response to environmental stimuli

Tissue microenvironment can influence DC subset distribution (*26, 27*). To examine how the major tuDC subsets identified in our FS tumors were distributed in other tissues, we performed scRNA-seq of Zbtb46-GFP^+^ DCs from an unrelated tumor type (murine flank B16 melanoma) and normal tissue (lungs). B16 melanoma tuDCs contained all major tuDC subsets, including clusters enriched for *Ly6a, Cxcl9* and *Cxcl10* expression that resembled IFN-DCs (Figures 3A and 3B). On the other hand, healthy lung DCs consisted of cDC1s, CD11b^+^ DC2s and mDCs but lacked IFN-DCs and had minimal *Ly6a, Cxcl9* and *Cxcl10* expression (Figures3C and 3D). Using gene signatures derived from the FS tuDC subsets, we identified and compared populations across datasets, observing high variability in the proportion of IFN-DCs with healthy lungs showing the lowest proportion (Figure 3E). In contrast, the proportion of mDCs was consistent across datasets (Figure 3E). We also corroborated these findings by FC where Zbtb46-GFP^+^ DCs in B16 melanomas comprised of a sizeable population of CCR7^-^Ly6A^hi^ IFN-DCs that were nearly absent in normal lungs (Figure 3F).

**Figure 3:**
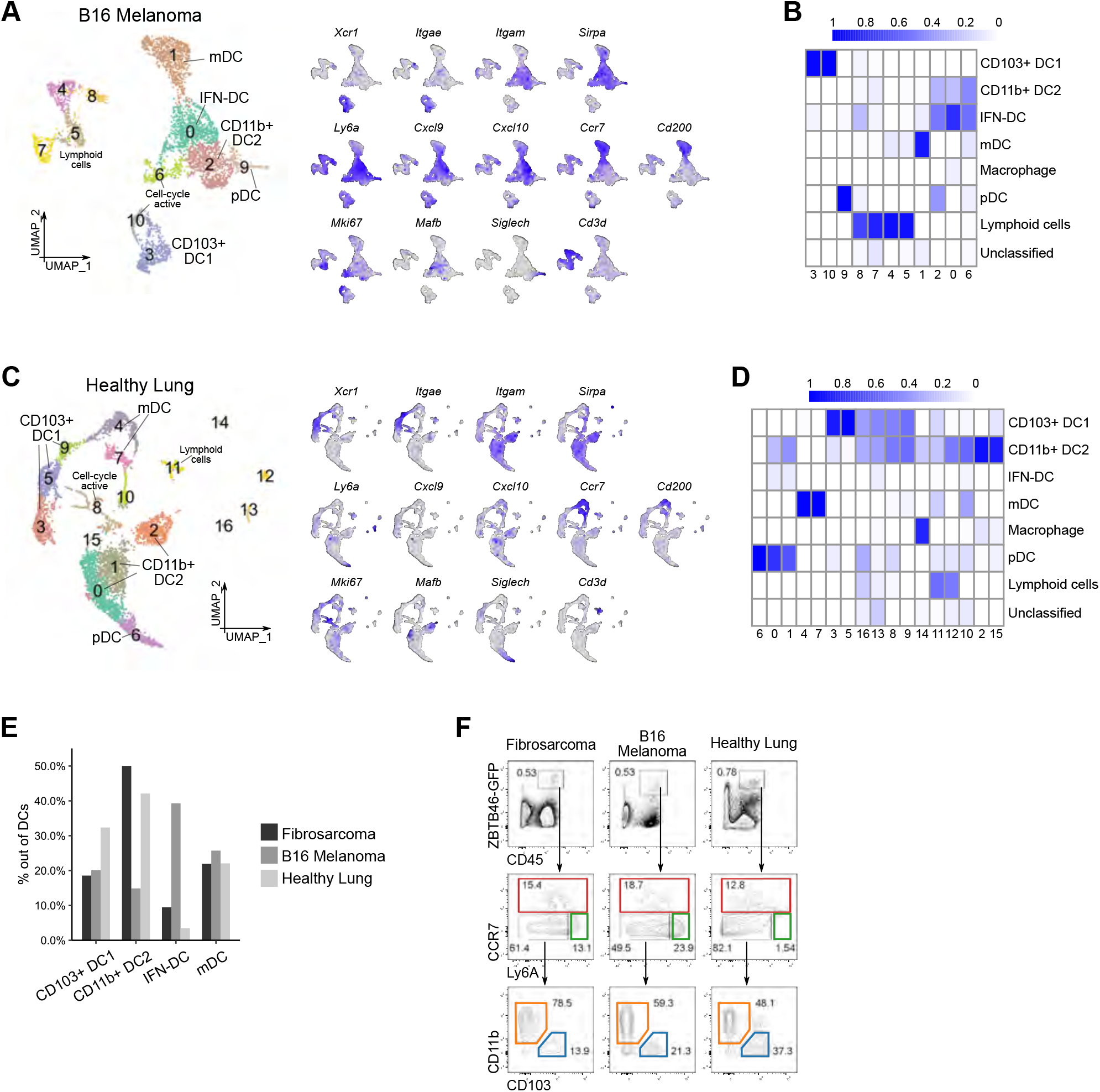
IFN-DC frequency differ between tissue and tumor. A) UMAP plot (left panel) of scRNA-seq data of 4,080 sorted live CD45+ZBTB46-GFP+ DCs from B16 melanoma. Shown are the numbered clusters identified by Seurat analyses and subset identities based on expression of marker genes. UMAP plots on the right show expression of the subset annotation markers (gray = low expression; blue = high expression) (right panel). B) Heatmap displaying annotation of cells in the B16 melanoma scRNA-seq dataset using reference gene expression data from the FS tuDC scRNA-seq. Scores represent the proportion of cells within a cluster assigned to a specific cell-type using *SingleR*. C-D) Shown are equivalents of (A) and (B) for scRNA-seq data of 5,224 sorted live CD45+ZBTB46-GFP+ DCs from lungs of wild type (C57BL/6J) mice. E) Relative proportions of the four major DC subsets (Y-axis) out of total DCs within each color-coded single-cell data sample. F) FC profiling of DC subsets in FS tumors, B16 tumors and healthy lung tissue in ZBTB46-GFP hosts.

A common method to generate cDCs *in vitro* involves bone marrow cells cultured in the presence of FLT3 ligand (FLT3L) (*28*). Zbtb46-GFP^+^ cells in these FLT3L driven bone marrow-derived DCs (F-BMDC) cultures also contained CCR7^-^Ly6A^hi^ and CCR7^+^ populations (Figure S3A). Thus, we next asked whether these populations resemble the *in vivo* tuDC subsets, in particularly IFN-DCs and mDCs, by performing scRNA-seq of bulk F-BMDC cultures (Figure S3B). Clusters consistent with *in vivo* cDC1s, CD11b^+^ DC2s and mDCs were identified in F-BMDCs (Figure S3C). However, in contrast to *in vivo* DCs, there was broad expression of *Ly6a* but absent expression of *Cxcl9* and *Cxcl10* in F-BMDCs (Figure S3B). Correspondingly, we were unable to find IFN-DCs in F-BMDCs (Figures S3B and S3C). Thus, although F-BMDCs consisted of CCR7^-^Ly6A^hi^ populations by FC, they did not resemble *in vivo* IFN-DCs.

Taken together, findings in this section suggested that IFN-DCs are seen in tumors but not normal lungs or *in vitro*, implying that tumor-associated factors may lead to their appearance.

### IFN-DCs originate from CD11b+ dendritic cells in response to interferons

A few recent studies have shown the presence of DCs enriched for IFN signaling reminiscent of our IFN-DCs (*29–31*). Bosteels and colleagues showed that in a murine model of viral respiratory infection, IFN-I acts on cDC2s to generate a DC subset (inf-cDC2s) characterized by an IFN signature that helps prime CD4^+^ and CD8^+^ T cell responses (Bosteels et al., 2020). More recently, Duong and colleagues described similar IFN-I-induced CD11b^+^ DCs that promoted anti-tumor immune responses through antigen cross-dressing (Duong et al., 2021). Finally, Ghislat and colleagues described the presence of DCs enriched for IFN signaling within the cDC1 subset (*30*). Thus, IFN-I signaling appears to regulate a specific subset of DCs in infection and cancer but how these DCs are different from other DC subsets and the role of type 2 interferons (IFN-II) in their generation remain to be fully elucidated. Hence, we further examined the regulation and origins of our IFN-DCs.

Analysis of IFN-DCs in the FS tuDC dataset at a higher clustering resolution revealed two further subsets, which we denoted as IFN1-DC and IFN2-DC (Figure 4A). Gene expression comparison between the two subsets indicated differential expression of interferon-stimulated genes (Figure 4B). For example, *Irf7* was upregulated in IFN1-DCs and *Cxcl9* was upregulated in IFN2-DCs. On the other hand, *Cxcl10* and *Ly6a*, our general marker of IFN-DCs, were equivalently expressed in both subsets. Further examination of interferon signatures revealed enrichment of IFNα and IFNγ signaling in IFN1-DCs and IFN2-DCs respectively (Figure 4C). IFN1-DCs were also enriched for the signature of ISG^+^ DCs – a recently described IFN-I-regulated tuDC subset (Duong et al., 2021) (Figure 4C). Correspondingly, the ISG^+^ DC marker *Axl* was higher in IFN1-DCs compared to IFN2-DCs (Figure 4B). Based on these observations, we were able to distinguish IFN1- and IFN2-DCs in FC by using Ly6A and AXL (Figure S4A). Of note, consistent with their sessile nature, both subsets were detected in the tumor but not in the tumor-draining lymph node (Figure S4B). Ingenuity Pathway Analysis of DEGs between the two IFN-DC subsets predicted top upstream regulators of IRF7 and IFNα for IFN1-DCs, and STAT1 and IFNγ for IFN2-DCs (Figure S4C). Thus, we tested the role of IFN-I and IFN-II in regulating IFN-DCs through *in vivo* treatments with neutralizing antibodies (αIFNAR1 or αIFNγ). As predicted, either treatment significantly reduced the frequency of CCR7^-^Ly6A^hi^ IFN-DCs within tuDCs (Figures 4D and S4D).

**Figure 4:**
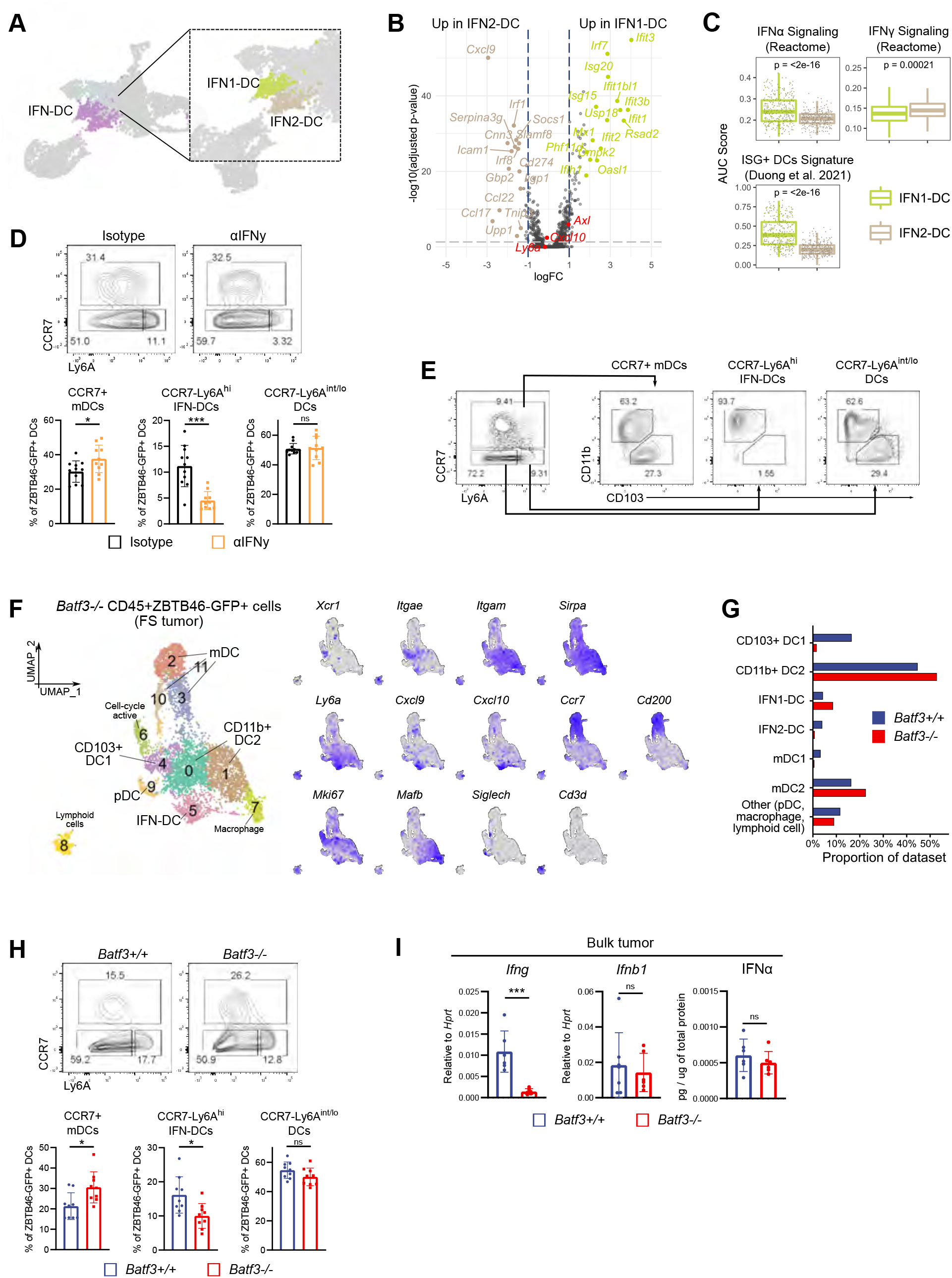
Interferons generate IFN1- and IFN2-DCs from CD11b+ DC2s. A) UMAP plot of the IFN-DCs from the FS scRNA-seq dataset re-clustered at a higher resolution showing IFN1-DC and IFN2-DC. B) Comparison of DEGs between IFN1-DCs and IFN2-DCs from the FS scRNA-seq. The top 15 DEGs for each subset along with selected genes of interest are highlighted. C) Boxplots comparing the relative enrichment of the Reactome IFNα and IFNγ signaling gene sets (top), and the ISG+ DC signature (from Duong *et al.* 2021) (bottom) between IFN1-DCs and IFN2-DCs in the FS scRNA-seq as calculated via *AUCell*. D) FC profiling of tuDCs from FS tumors treated with control isotype or αIFNγ blocking antibody (200 ug intraperitoneal on days 4, 7, 10 and 13 post-tumor injection). Representative FC plots pre-gated on live CD45+ZBTB46-GFP+ tuDCs (upper panel). Frequency of tuDC subsets within total tuDCs pooled from two independent experiments (lower panel). E) FC profiling of tuDCs from FS tumors for CD11B and CD103 expression. Events are pre-gated on live CD45+ZBTB46-GFP+ tuDCs. F) UMAP plot of scRNA-seq data of 6,456 sorted live CD45+ZBTB46-GFP+ cells from FS tumors in *Batf3^-/-^ Zbtb46^GFP^* mice (left panel). Shown are the numbered clusters identified by Seurat analyses and assigned subset identities based on expression of marker genes. UMAP plots on the right show expression of subset annotation markers (gray = low expression; blue = high expression). G) Relative proportions of the tuDC subsets within the scRNA-seq of FS tumor DCs in *Batf3^+/+^* ZBTB46^GFP^ and *Batf3^-/-^* ZBTB46^GFP^ mice. H) FC profiling of tuDCs from FS tumors in *Batf3^+/+^*ZBTB46^GFP^ and *Batf3^-/-^* ZBTB46^GFP^ hosts. Representative FC plots pre-gated on live CD45+ZBTB46-GFP+ tuDCs (upper panel). Frequency of tuDC subsets within total tuDCs pooled from two independent experiments (lower panel). I) Levels of interferons within FS tumors in *Batf3^+/+^* and *Batf3^-/-^* hosts as measured by RT-qPCR (*Ifng* and *Ifnb1*) or by ELISA (IFNα). \**p* < 0.05, \*\**p* < 0.01, \*\*\**p* < 0.001. Statistical analysis was performed with unpaired *t* test (C, D, H, and I). Data are shown as mean ± SD.

We next examined the ontogeny of IFN-DCs. As described above (Figure 1C), mDCs comprised of mDC1s and mDC2s based on their gene expression pattern. Supporting this, FC showed discrete CD11b^+^CD103^-^ and CD11b^-^CD103^+^ populations within CCR7+ mDCs (Figure 4E). In contrast, IFN-DCs were entirely made up of CD11b^+^CD103^-^ cells (Figure 4E). This observation combined with the above described (Figure 1D) transcriptional similarity of IFN-DCs with CD11b^+^ DC2s indicated that IFN-DCs likely originate from CD11b^+^ DC2s. To rule out a cDC1 origin for IFN-DCs more definitively, we examined *Batf3*^-/-^ mice that lack cDC1s (*9*). We generated *Zbtb46*^GFP/+^: *Batf3^-/-^* mice and isolated Zbtb46-GFP^+^ tuDCs from FS tumors in these mice for scRNA-seq analysis. In wild-type (*Batf3^+/+^*) tuDCs, *Batf3* was highly expressed in cDC1s and mDCs (Figure S4E). *Batf3^-/-^*tuDCs lacked both resting and migratory cDC1s but maintained resting and migratory CD11b^+^ DC2s (Figures 4F and 4G). Surprisingly, IFN2-DCs, but not IFN1-DCs, were selectively reduced in *Batf3*^-/-^ tuDCs, which was also supported by a general reduction in CCR7^-^Ly6A^hi^ IFN-DCs by FC analysis (Figures 4G and 4H). Correspondingly, sorted CCR7^-^CD11b^+^ tuDCs showed a significant reduction in the expression of IFN2-DC-enriched genes (*Cxcl9* and *Gbp2*), a small reduction in the expression of genes associated with both IFN-DC subsets (*Cxcl10*), and no changes in the expression of IFN1-DC-enriched genes (*Irf7*) (Figure S4F). This may indicate a cell-intrinsic (e.g., cDC1 origin) or a cell-extrinsic requirement of *Batf3* for IFN2-DCs. Given the role of IFNs in regulating the two IFN-DC subsets, we hypothesized that it could be the latter scenario and evaluated the levels of IFNs in the TME (Figure 4I). IFN-I (both *Ifnb1* and IFNα) were unchanged between wild-type and *Batf3*^-/-^ but *Ifng* was significantly decreased in *Batf3^-/-^* tumors, which can explain the reduced IFN2-DCs in *Batf3*^-/-^.

We next explored sources of IFNγ for the induction of IFN2-DCs. T cells and NK cells produced the most *Ifng* in the TME of wild-type FS tumors (Figure S4G). *Batf3*^-/-^ mice lack cDC1s, which results in the selective loss of cytotoxic CD8+ T cells (CTLs) in the TME (Figure S4H) (*9*). The reduction of CD8^+^ T cells can explain decreased *Ifng* and attendant loss in IFN2-DCs in *Batf3^-/-^* tumors. Further supporting this notion, depletion of T cells led to significant reductions in CCR7^+^ mDCs and CCR7^-^Ly6A^hi^ IFN-DCs in FS tumors (Figure S4I). These findings underscored the importance of cross talk between CTLs and DCs (via IFNγ) in the TME.

Immunotherapy with checkpoint blockade work by reinvigorating and enhancing T cell functions in tumors (*32*), and thus we next examined potential correlations between IFN-DCs and IFNγ in the αPD1 therapeutic setting. Treatment of FS tumors with αPD1 increased *Ifng* levels and IFN-DCs in the TME but not mDCs (Figure S4J and S4K). Responses to αPD1, as measured through tumor size differences, were variable in this FS tumor model. Nonetheless, within the αPD1-treated group we observed a significant negative correlation between tumor growth and intratumoral CD8^+^ T cell frequency (Figure S4L). Notably, tumor growth also had negative correlation trends with frequencies of IFN-DCs and mDCs, with IFN-DCs showing a stronger negative correlation (Figure S4L). We also found a small trend of positive correlations between IFN-DC frequency with *Ifng* levels and intratumoral CD8^+^ T cell frequency. Collectively, these findings indicated that IFN-DCs were regulated by local environmental IFN levels, which involved T cell-mediated IFNγ in FS tumors.

### IFN-DCs are present in human tissue

Major DC subsets have shown strong conservation between mice and humans at the transcriptional and functional level in different contexts including in tumors (*21, 26, 33*). We asked whether insights from our murine studies on IFN-DCs could be extended to scRNA-seq of human MNPs. We utilized a recently published comprehensive database of human MNPs (MNP-VERSE) curated from 41 different scRNA-seq datasets (*34*). The datasets represented 13 tissue types containing both normal and diseased tissues including cancer. We extracted and re-analyzed cells from the MNP-VERSE that the authors had annotated as DCs from healthy and cancer tissues (Figure S5A). Unsupervised graph-based clustering of these pre-annotated DCs identified cells corresponding to the major DC subsets in our murine studies: DC1s (cluster 2; *XCR1* and *CLEC9A* expression), DC2s (clusters 0, 3 and 6; *CD1C* and *FCER1A* expression), IFN-DCs (cluster 10; *CXCL9* and *CXCL10* expression), and mDCs (cluster 5; *CCR7* and *LAMP3* expression) (Figure 5A). Beyond the major DC subsets, we also identified clusters representing macrophages (Mac; cluster 1; *APOE* expression), monocytes (Mono; cluster 4; *S100A12* expression), proliferating DCs (cluster 9; *MKI67* expression), and Langerhan cells as well as cells with gene expression profiles not associated with cDC identity (Other; clusters 7, 8, 11-14; *CD207*, *CD3D*, *GZMB* and *LTB* expression) (Figures 5A and S5B). As expected, the relative frequency of these subsets differed between the 31 datasets used as the source of pre-annotated DCs (Figure S5C). Like murine mDCs (Figures 1E and 2A), human mDCs displayed highest enrichment for the ‘DC Maturation ON’ signature followed by IFN-DCs, while the IFN signaling signatures (both IFNα and IFNγ) were most enriched in IFN-DCs (Figure 5B).

**Figure 5:**
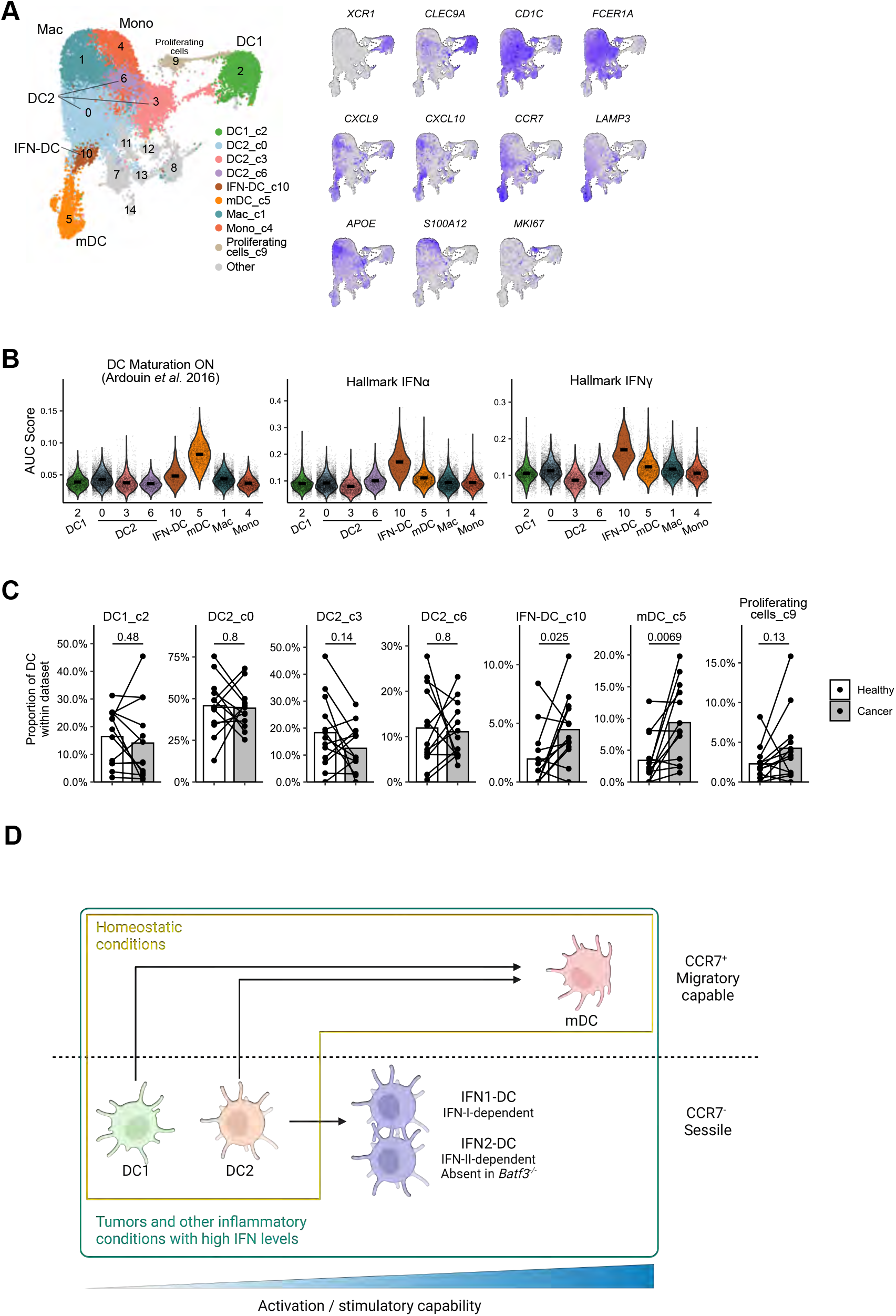
Human tumors and normal tissue contain murine equivalents of IFN-DCs. A) UMAP plot of pre-annotated human DCs from healthy and cancer tissues of the MNP-verse (Mulder et al., 2021) (left panel). Numbered clusters and assigned subset identities based on expression of marker genes are shown. UMAP plots on the right show expression of subset annotation markers (gray = low expression; blue = high expression) (right panel). B) Violin plots comparing the relative enrichment of DC maturation and interferon response hallmark gene sets among defined clusters as calculated via *AUCell*. Crossbar within violin represents mean. C) Relative proportions of DC subsets among total DCs from paired healthy or cancer tissues. Quantification was restricted to non-lymphoid tissue datasets that contained cells of both healthy or cancer status (*n* = 13). Statistical analysis was performed with paired *t* test. Horizontal line height represents mean. D) Model based on our findings. Tissue DCs mature and migrate to draining lymph nodes under homeostatic conditions. During inflammation, high levels of local interferons generate IFN-DCs from CD11b+ DC2s (CDC2 or MoDC) that are activated but largely sessile. In contrast to migratory DCs that drive T cell priming, we posit that IFN-DCs drive T cell infiltration and stimulation in the local microenvironment.

We next compared human DC subsets between healthy and cancer tissues. For consistency, we restricted our analysis to DCs from datasets of non-lymphoid tissues that contained cells of both healthy and cancer status. (Figure S5A). The proportions of IFN-DCs and mDCs within total DCs were higher in cancer tissues compared to that of healthy tissues (Figure 5C). This is consistent with the notion that IFN-DCs and mDCs are two activated DC subsets and hence increased in inflammatory conditions such as tumors.

In closing, our findings clarified DC heterogeneity in tumors and identified IFN-DCs as an activated but sessile DC subset that originated from CD11b^+^ DCs in the presence of high local IFN levels. Based on our findings, we propose a dynamic model intratumoral DC composition that co-evolves with local adaptive immune response (Figure 5D).

## DISCUSSION

Single cell genomics is providing unprecedented insights into immune cell heterogeneity. This approach is relatively straightforward for abundant immune cells but challenging for rare populations, especially in inflammatory conditions where cells often display overlapping expression of surface markers. Our data demonstrates the advantage of using genetic reporter mice (*Zbtb46*^GFP/+^) to rapidly identify and isolate rare immune cells for scRNA-seq. Given the higher purity and numbers of DCs compared to previous studies, we were able to uncover novel DC states. This work also establishes scRNA-seq profiles of Zbtb46-GFP^+^ DCs from (1) lungs, (2) fibrosarcomas, (3) melanoma, and (4) fibrosarcomas generated in *Batf3^-/-^* mice, as well as DCs generated *in vitro* with the FLT3L culture system, which will be valuable to the larger immunology community.

A few recent studies have also mentioned cells reminiscent of our IFN-DCs (*29–31, 35*). Given the overlapping observations and the emerging status of this DC subset, we highlight key similarities and differences between our findings with these recent studies. Firstly, IFN-DC-like cells generated in response to IFN-I have been described (*29, 31*). In our study, we find that IFN-DCs comprised of two subsets – one with an IFN-I signature (IFN1-DC), and therefore similar to previously characterized subsets, and the other with an IFN-II signature (IFN2-DC). Indeed, we can distinguish these subsets of IFN-DCs in FC by a combination of Ly6A (as a marker of overall IFN-DCs) and AXL (as a marker of IFN1-DC). While Duong and colleagues observed tumor-intrinsic IFN-I as the source of interferon for their ISG^+^ DCs (*31*), our studies show a role of T cell-derived IFNγ in generating IFN2-DCs. Thus, our work complements existing studies and together highlights the relevance of this emerging DC subset, although further studies are required to dissect the precise roles of IFN1-DCs vs IFN2-DCs in tumors. Secondly, IFN-DC-like cells were characterized as migratory DC in a murine respiratory infection model (*29*), or were characterized as a transitionary state to migratory DCs in tumors based on *in silico* analyses (*30, 35*). In contrast, our study identifies IFN-DCs as CCR7^-^ cells and we show that key markers of IFN-DCs (e.g. Ly6A, CXCL9, and AXL) were not expressed among migratory DCs in the tumor draining lymph nodes. Additionally, the inability of sorted tumor IFN-DCs to significantly upregulate *Ccr7* or migrate towards CCL19 over time compared against other CCR7^-^ subsets suggests that IFN-DCs have significantly lower migratory potential or were largely sessile. Moreover, as was previously described (*29, 31*), it is noteworthy that IFN-DCs acquire expression of markers traditionally associated with macrophages (e.g. CD64 (*Fcgr1*) and AXL), an antigen-presenting cell that is known to be sessile (*36, 37*).

Our findings highlight IFN-DCs and mDCs as two activated DC states with discrete properties. CXCL9, a key driver of T cell infiltration, is highly expressed by IFN-DCs (specifically IFN2-DCs) but has very limited expression in CCR7^+^ mDCs. Previous studies have proposed cDC1s to be the primary source of CXCL9 among tuDCs (*38–40*). We also find that cDC1s can produce CXCL9, although at lower numbers than that of IFN-DCs. Moreover, our identification of IFN-DCs as a major source of CXCL9 is likely aided by our use of *Zbtb46*^GFP/+^, which enables clear distinction of *bona fide* DC2s from tumor macrophages that are also known to produce CXCL9 (*41*). Thus, while T cells following a CXCL9 gradient can ‘meet’ a naïve cDC1, macrophage, or IFN-DC in tumors, IFN-DCs are superior in their ability to activate or re-stimulate these T cells.

Successful CTL-dependent anti-tumor responses involve re-activating CTLs via interactions with stimulatory cells in the TME (*8, 40*), though the nature of these stimulatory cells remains under investigation. mDCs have been implicated in this role (*20, 42*), with one recent publication showing clusters of mDCs in tumor stromal perivascular niches that provide pro-survival signals to infiltrating CTLs (*43*). Our findings suggest that IFN-DCs are also suited to act as intratumoral re-activators of CTLs due to their sessile nature, their expression of T cell chemoattractants, and their ability to cross-dress or cross-present antigen and activate CD8+ T cells (*29, 31*). Thus, we propose a dynamic model of DCs in tumors that includes spatiotemporal division of labor between activated DCs (Figure 5D). At homeostasis, DCs mature and migrate to draining lymph nodes. If adaptive immune responses (against tumors as an example) are induced by these mDCs, the ensuing influx of CTLs results in increased IFNγ levels in the TME. This leads to generation of IFN2-DCs, which along with IFN1-DCs, attract and re-activate CTLs intratumorally for continued immune responses against tumors. In this context, our findings regarding changes in IFN-DCs with immune checkpoint blockade is intriguing and important as they point towards a potential role of this DC subset in mediating or prognosticating responses to immunotherapy, although further work is needed to explore this link

## MATERIALS AND METHODS

### Animals

*Zbtb46*^GFP^ mice were a generous gift from Dr. Kenneth Murphy. *Lyz2*^Cre^ mice and *Rosa26*^LSL-tdT^ mice were obtained from Jackson Laboratories and bred together with *Zbtb46*^GFP^ mice to generate *Zbtb46*^GFP^ *Lyz2*^Cre^ *Rosa26*^LSL-tdT^ mice. *Batf3^−/−^* mice were obtained from Jackson Laboratories and bred to *Zbtb46*^GFP^ mice to generate *Zbtb46*^GFP^ *Batf3^−/−^* mice. C57BL/6 and OT-I mice were obtained from Jackson Laboratories.

Mice of either sex ranging in age from 8-12 weeks were used for these studies. Mice were bred and maintained in specific pathogen free facilities at the University of Pennsylvania. All animal procedures were conducted according to National Institutes of Health guidelines and approved by the Institutional Animal Care and Use Committee at the University of Pennsylvania.

### Tumor cell lines, implantation of tumor cells, and tumor growth measurements

The C57BL/6 syngeneic fibrosarcoma cell line (FS) was a generous gift from Dr. Robert Schreiber and the B16-F10 melanoma cell line (B16) was a generous gift from Dr. Andy Minn. Tumor cell lines were cultured in complete DMEM media (DMEM media containing 10% FBS, 1% Pen/Strep and 2 mM glutamine). For implantation of tumor cells, low passage (< P15) cell lines were detached using 0.05% trypsin (GIBCO) and washed once with complete DMEM media. Tumor cells were resuspended in PBS and implanted subcutaneously (s.c.) at 1×10^6^ in 200 µL or implanted intramuscularly (i.m.) into posterior thigh musculature (hip extensors) at 1×10^6^ in 50 µL. B16 cells were implanted s.c. for all experiments involving this tumor. FS cells were implanted s.c. for the generation of the scRNA-seq datasets and were implanted i.m. for all other experiments involving FS samples. Growth of FS i.m. tumors were determined by taking the difference between the diameter of the tumor-bearing hind leg at tumor implantation and harvest.

### *In vivo* treatments

For blocking IFN signaling, mice were treated i.p. with 200 µg of rat IgG1 isotype control (clone HRPN), αIFNAR1 for IFN-I blockade (clone MAR1-5A3) or αIFNγ for IFN-II blockade (clone XMG1.2) at days 4, 7, 10 and 13 post-tumor implantation. For immune checkpoint blockade, mice were treated i.p. with 200 µg of rat IgG2a isotype control (clone 2A3) or αPD1 (clone RMP1-14) at days 6, 9 and 12 post-tumor implantation. For depletion of T cells, mice were treated i.p. with buffer alone or 200 µg αCD3 (clone 17A2) at days -3, 0, 3, 6 and 9 post-tumor implantation. All *in vivo* antibodies were purchased from BioXCell.

### Tissue harvest and processing

All tumors were harvested at day 14 post-tumor implantation unless otherwise stated. Tissues were chopped with surgical scissors and then digested in 30 U/mL of DNase I (Sigma-Aldrich) and 250 µg/mL of Collagenase B (Roche) to prepare single cell suspensions. Tumors and lungs were digested at 37°C for 40 minutes with stirring, and spleens and lymph nodes were digested at 37°C for 25 minutes without stirring. Cell suspensions were then filtered twice through a 70 µm cell strainer. A red blood cell (RBC) lysis step using 1X RBC Lysis Buffer (BioLegend) was performed in between the two filtrations for samples from tumors, lungs and spleens.

### Flow cytometry and cell sorting

Single cell suspensions were first incubated for 10 minutes on ice with anti-mouse CD16/32 Fc Block (BD Biosciences), and subsequently stained with fluorophore-conjugated antibodies against surface markers at 4°C for 20 mins. CCR7 staining was performed separately at 37°C for 30 mins prior to staining for other markers. For CXCL9 staining, single cell suspensions were incubated with 5 µg/mL of brefeldin A at 37°C, 5% CO_2_ for 5.5 hours, stained for surface markers, and then stained for intracellular CXCL9 using a fixation/permeabilization kit (BD Biosciences) according to manufacturer’s instructions.

Sample acquisition was performed on a LSRFortessa flow cytometer (BD Biosciences) and analyzed using FlowJo software (Tree Star, Inc.). When applicable, absolute cell numbers were quantified using 5 µL of 123count eBeads Counting Beads (Thermo Fisher Scientific) added to each sample.

For sorting of tumor APC populations, samples were first positively selected for CD11c^+^ cells using Mouse UltraPure CD11c Microbeads (Miltenyi Biotec) prior to surface marker staining. Cell sorting was performed on a FACSAria II (BD Biosciences) and cells were sorted into chilled complete IMDM media (10% FBS, 1% Pen/Strep, 2 mM glutamine, 0.05 mM 2-ME, 1X NEAA and 1 mM NaPyr).

### APC-T cell co-cultures

For co-cultures with T cells activated by αCD3/CD28 beads, pan splenic T cells were isolated from crushed spleens of naïve C57BL/6 mouse using the Mouse Pan T Cell Isolation Kit (Miltenyi Biotec). In the subsequent co-culture incubation, T cells were activated by 2 µL of Mouse αCD3/28 Dynabeads (Thermo Fisher Scientific).

For co-cultures with CD8^+^ OT-I T cells activated by SIINFEKL presentation, CD8^+^ OT-I splenic T cells were isolated from crushed spleens of naïve OT-I mice using the Mouse CD8 T cell Isolation Kit (BioLegend). Sorted tumor APC populations were pulsed with 0.01 ng/mL SIINFEKL peptide (Invivogen) at 37°C for 1.25 hours and then washed twice.

For all APC-T cell co-cultures, T cells were labeled with 2.5 µM of CellTrace CFSE dye (Thermo Fisher Scientific) at room temperature for 5 minutes before washing twice. 5×10^3^ APCs were co-cultured with 2.5×10^4^ CFSE-labeled T cells in 200 µL of RPMI media containing 10% FBS, 1% Pen/Strep, 2 mM glutamine, 20 mM HEPES, 1 mM NaPyr and 0.05 mM 2-ME. After 3 days of incubation at 37°C, T cell proliferation was measured by CFSE dilution and CD44 expression via flow cytometry.

### APC transwell migration assay

After sorting, 1.5-5.0×10^4^ tumor APC populations were added to the upper chambers of transwells (6.5 mm, 5.0 µm Pore Polycarbonate Membrane Insert; Corning) in 100 µL of complete IMDM media (IMDM media containing 10% FBS, 1% Pen/Strep, 1% glutamine, 1% NEAA, 1 mM NaPyr and 0.05 mM 2-ME). The lower compartments were filled with 600 µL of complete IMDM media with or without 200 ng/mL of CCL19 (PeproTech). After incubation at 37°C for 21 hours, cells in the lower chambers were harvested and counted using flow cytometry. For each APC population, the relative specific migration was calculated as follows: (number of cells in the lower chamber with CCL19 – number of cells in the lower chamber without CCL19) / total inputted cells to the upper chamber.

### *Ccr7* and *Cxcl9* induction assay

After sorting, 1×10^4^ specified tumor APC populations were added to a flat-bottom 96-well plate. One set of samples was immediately harvested to serve as a baseline control, while remaining samples were resuspended in 200 µL of complete DMEM media or tumor-conditioned media (TCM). TCM was prepared by filtering supernatant from 2 day old sub-confluent FS cell line cultures that were seeded at 1-1.5×10^6^ in 10 mL of complete DMEM media per 10 cm plate. Following 20 hrs of incubation at 37°C and 5% CO_2_, samples were spun down and lysed with TRIzol (Thermo Fisher Scientific) for subsequent RNA extraction.

### FLT3 ligand bone marrow cultures

Single bone marrow cell suspensions were plated at 4×10^6^ per well in 2 mL of complete IMDM media supplemented with 150 ng/mL of FLT3L (PeproTech). On day 7 of differentiation, floating cells were harvested for analysis.

### RNA isolation and qPCR analysis for gene expression

To analyze gene expression of sorted APCs, 1-2×10^4^ cells were lysed in 200 µL of TRIzol (Thermo Fisher Scientific) and RNA was isolated according to manufacturer’s instructions. To analyze gene expression of bulk tumor samples, 100-150 mg of snap-frozen tissue were mechanically homogenized in 750 µL of Lysis Solution/2-mercaptoethanol mixture and total RNA was isolated using the GenElute Mammalian Total RNA Miniprep Kit (Sigma-Aldrich). Reverse transcription for up to 2 µg of total RNA was performed using the High-Capacity RNA-to-cDNA Kit (Thermo Fisher Scientific). qPCR was performed in a ViiA 7 Real-Time PCR machine (Thermo Fisher Scientific) using TaqMan Universal PCR Master Mix and probes (Thermo Fisher Scientific) for gene specific amplification. Relative gene expression levels were calculated as follows: 2^−(CT_gene of interest_−CT_*Hpart*_^).

### ELISA

Snap-frozen tumor samples were mechanically homogenized in 250 µL of 1X RIPA buffer (Cell Signaling Technologies) with 1X Halt protease inhibitors (Thermo Fisher Scientific) for every 100 mg of tissue. From the resulting lysate supernatant, total protein concentration was quantified using the Pierce BCA kit (Thermo Fisher Scientific) and IFNα concentration was quantified using the High Sensitivity Mouse IFNα All Subtype ELISA Kit (PBL Assay Science) according to the manufacturers’ instructions.

### Single Cell RNA Sequencing

#### Sample preparation, scRNA-sequencing and pre-processing

Mouse tissue samples for scRNA-seq were first positively selected for CD45^+^ cells using Mouse CD45 Microbeads (Miltenyi Biotec) and then sorted for 7AAD^-^CD45^+^Zbtb46-GFP^+^ expression before submission. F-BMDCs for scRNA-seq were submitted without prior enrichment. Preparation of sequencing libraries and sequencing were conducted at the Center for Applied Genomics Core at the Children’s Hospital of Philadelphia. Pooled libraries were prepared using the 10x Genomics Chromium Single Cell 3′ Reagent kit v3 per manufacturer’s instructions. Sequencing was performed on either an Illumina HiSeq 2500 or NovaSeq 6000. Demultiplexing and alignment of sequencing reads to the mm10 transcriptome and creation of feature-barcode matrices was done using the CellRanger pipeline (10X Genomics, v.3.0.0/3.0.2/3.1.0/6.0.0).

#### Standard scRNA-seq analysis

For each sample, the processed output files from CellRanger (barcodes, genes, matrix) were passed to the *Seurat* package (v.4.0.4) for downstream processing in *R* (v.4.1.1). Genes expressed in less than 3 cells were filtered out. Outlier cells were identified based on library size, number of expressed genes and mitochondrial proportion using *quickPerCellQC* in the *scater* package (v.1.20.1) and were subsequently removed. Normalization by deconvolution was performed using *computeSumFactors* in the *scran* package (v.1.20.1). The *Seurat* pipeline was used to identify highly variable genes (*FindVariableFeatures)* and principal component (PC) analysis was performed on the scaled 2000 most variable genes (*RunPCA*). The first 20 PCs were used for graph-based cluster identification (*FindNeighbors, FindClusters*) and UMAP dimensional reduction (*RunUMAP*). Clustering granularity was set at 0.5 resolution for clusters shown in Figures 1, S1, 3, S3, 4F and 5, and at 1.0 resolution for the separation of IFN1-DC and IFN2-DC in Figure 4A. Differentially expressed genes (DEGs) were identified using the default Wilcoxon rank sum test in the *FindAllMarkers* and *FindMarkers* functions from *Seurat*.

#### Annotation of cell identities by SingleR

Annotation of cell identities based on reference data was performed by *SingleR* (v.1.6.1). To annotate the FS WT tuDC dataset (Fig. 1D), cell identities were assigned using ImmGen reference data. Normalized expression data of mouse immune cell types from Phase 1 of the Immunological Genome Project (ImmGen) were downloaded via *celldex* (v.1.2.0). To annotate the B16, naïve lung, F-BMDC and FS *Batf3^-/-^* DC datasets (Figures 3B, 3D, 3E, S3C and 4G), the annotated FS WT tuDC scRNA-seq dataset was used as the reference data.

#### Relative enrichment of gene sets by AUCell

The interferon gene sets were obtained from hallmark v.7.0 and reactome v.7.2 on MSigDB. The DC maturation signatures were obtained from gene expression microarray comparisons of murine DCs matured under different conditions (Ardouin et al., 2016). To determine the relative enrichment of gene sets among cells, *AUCell* (v.1.14.0) was used to calculate an “Area Under the Curve” (AUC) score for each gene set in each cell.

#### Ingenuity Pathway Analysis for predicted upstream regulators

Statistically significant DEGs (< 0.05 adjusted *p* value) with greater than 0.5 log fold change between IFN1-DCs and IFN2-DCs in the FS WT tuDC scRNA-seq dataset, together with their respective fold change and adjusted *p* values, were supplied to Ingenuity Pathway Analysis (IPA) software. IPA returned the *p* value and activation z-score of predicted upstream regulators, of which the top 8 (ranked by *p* value) regulators with positive z-scores are shown for each subset in Figure S4C.

#### Analysis of human DCs

The entire MNP-VERSE was downloaded (https://gustaveroussy.github.io/FG-Lab/) (Mulder et al., 2021) and cells pre-annotated in the metadata as DCs (cDC1, DC2/DC3, mregDC) from healthy and cancer tissues were extracted out for further analysis. Within the extracted healthy and cancer DCs, datasets that contributed fewer than 100 cells and genes expressed in fewer than 100 cells were filtered out, resulting in a final analysis object of 25,238 pre-annotated DCs across 31 datasets. Integration across the 31 remaining datasets was performed using the *Seurat* reciprocal PCA workflow (*FindIntegrationAnchors, IntegrateData (*using 50 dimensions and 85 k.weights)). Following integration, unsupervised graph-based clustering and UMAP dimensional reduction were conducted as described above.

### Statistical Analysis

Statistical analyses were performed using GraphPad Prism 9 (GraphPad) or with the *R* package *ggpubr* (v.0.4.0). T test was used for comparison of two groups and one-way ANOVA with Tukey’s HSD post-test was used for multiple comparisons. *p* < 0.05 was considered statistically significant (\**p* < 0.05, \*\**p* < 0.01, \*\*\**p* < 0.001, \*\*\*\**p* < 0.001) and error bars represent SD.

## Supplementary Materials

**Figure S1:**
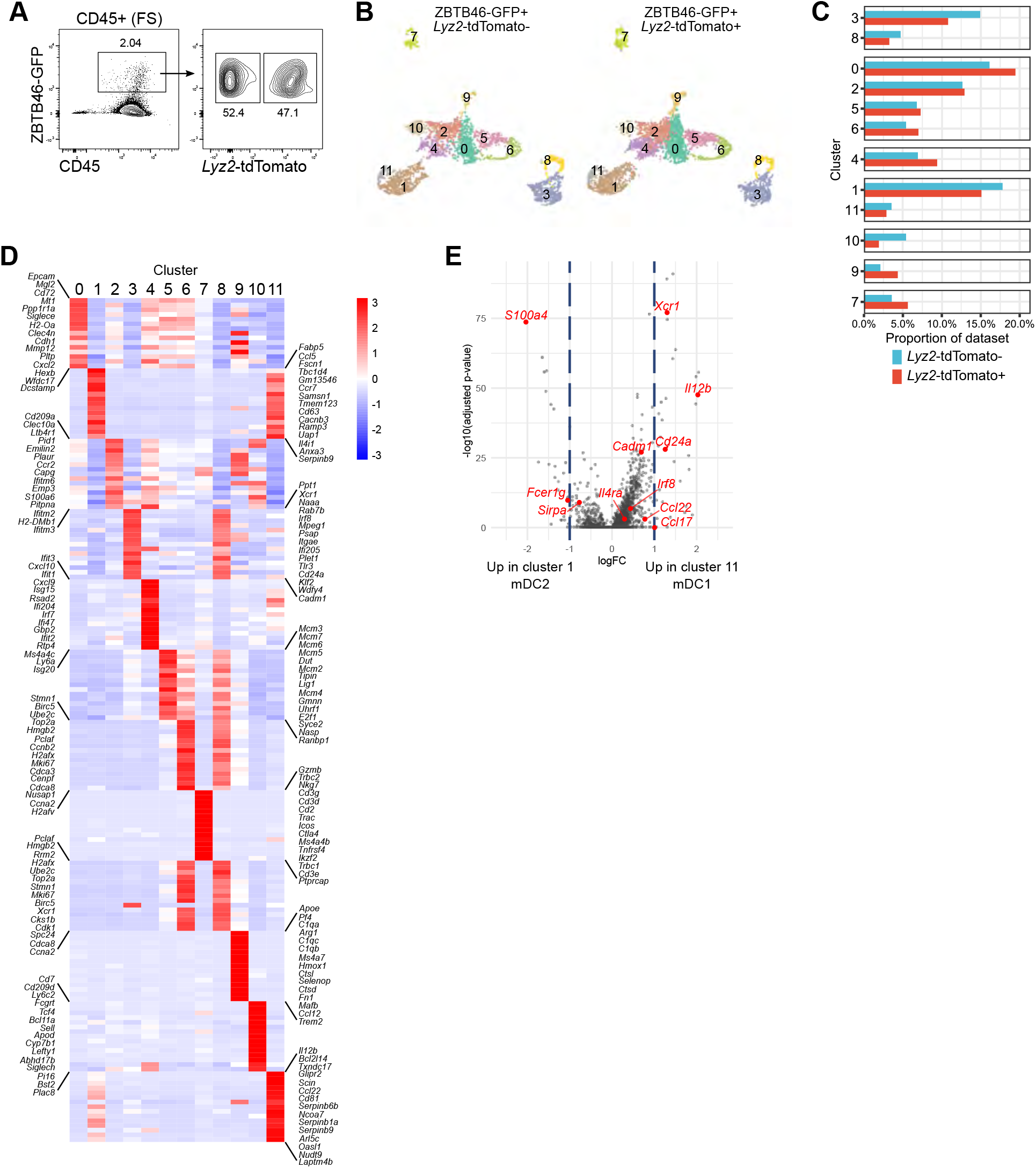
DC heterogeneity in solid tumors. Related to Figure 1. A) Gating strategy to sort *Lyz2*-tdT+ or *Lyz2*-tdT-cells within CD45+ZBTB46-GFP+ DCs from FS tumors for scRNA-seq (10x genomics). Single cell suspension from FS tumors were enriched for CD45+ cells using the Miltenyi^TM^ magnetic columns prior to cell sorting. B) Split UMAP plots of *Lyz2*-tdT+ and *Lyz2*-tdT-cells from CD45+ZBTB46-GFP+ DCs. C) Comparison of the distribution of clusters between *Lyz2*-tdT- and *Lyz2*-tdT+ subsets of CD45+ZBTB46-GFP+ DCs. D) Heatmap displaying top 15 marker genes per cluster (see Fig 1C). Shown are scaled, averaged normalized expression values for each cluster. E) Comparison of DEGs between the two mDC clusters, mDC2 (c1) and mDC1 (c11), from the FS scRNA-seq dataset with selected genes highlighted in red.

**Figure S2:**
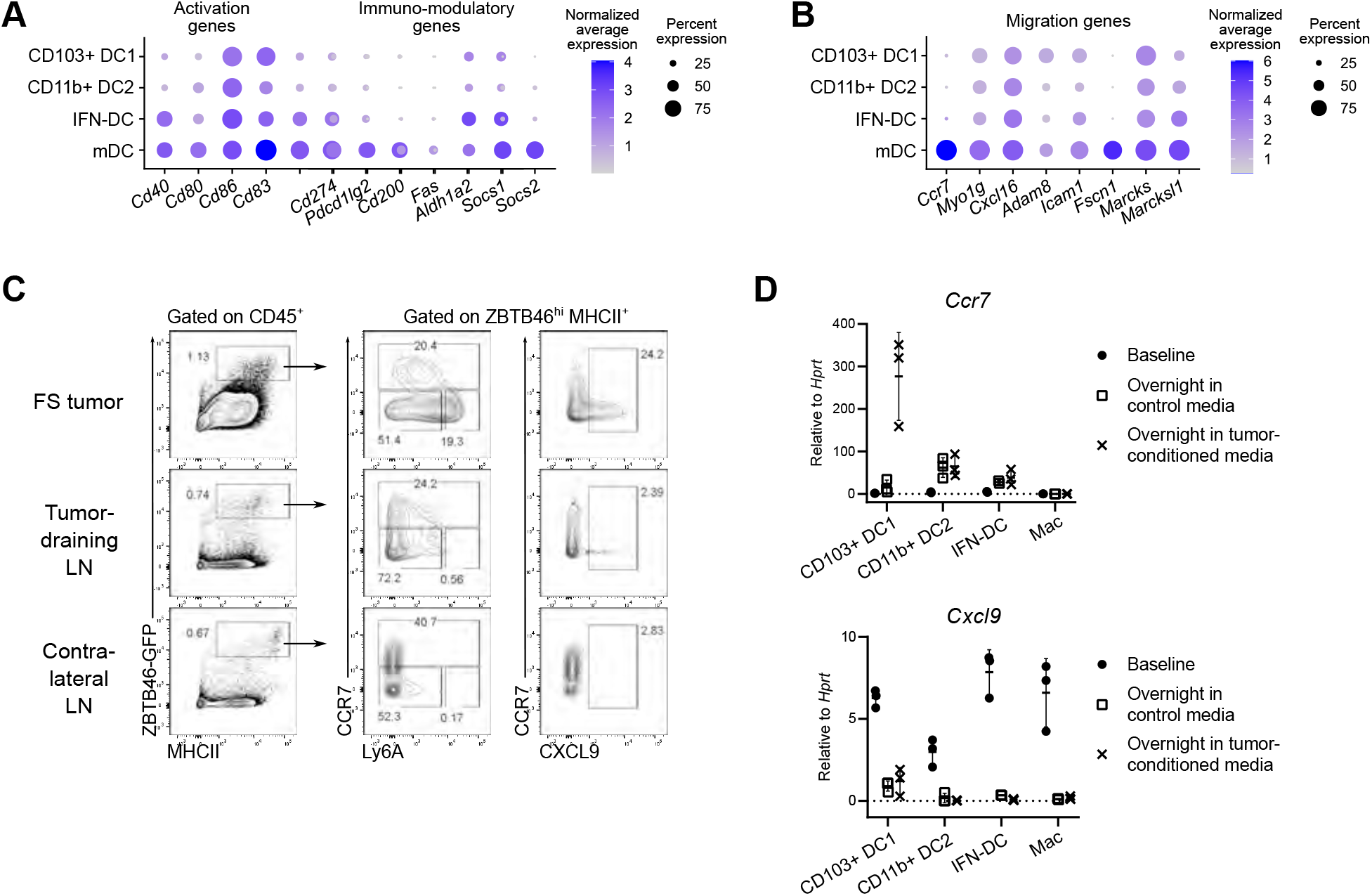
IFN-DCs and mDCs display distinct properties. Related to Figure 2. A) Dot plot showing expression of activation and immuno-modulatory genes (from Maier *et al.* 2020) among the major tuDC subsets in the FS scRNA-seq dataset. B) Dot plot showing expression of migration genes (from Maier *et al.* 2020) among major tuDC subsets in the FS scRNA-seq dataset. C) FC profiling of ZBTB46-GFP+ DCs with indicated markers in matched FS tumor, tumor-draining lymph node (tdLN) and contralateral (non tumor-draining) lymph node (cLN). D) Expression of *Ccr7* and *Cxcl9* (RT-qPCR) in sorted CCR7-tuDC subsets and macrophages at 0 h post-sorting (baseline) and after 20 h incubation in control or tumor-conditioned media. Data are shown as mean ± SD.

**Figure S3:**
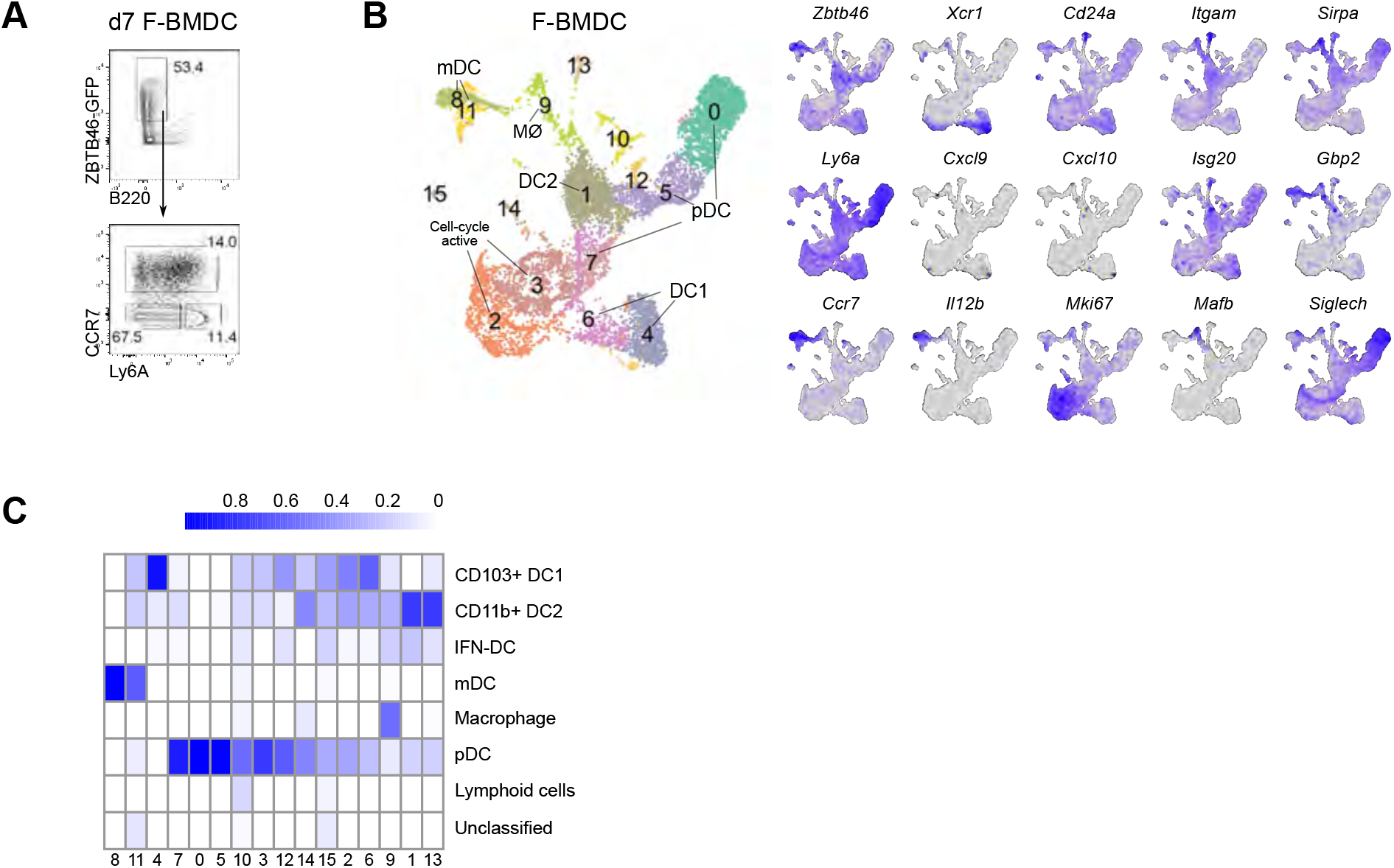
IFN-DCs are absent from DCs generated in vitro. Related to Figure 3. A) FC profiling of day 7 Flt3L bone marrow dendritic cells (F-BMDCs) generated from wild-type mice. B) UMAP plot of scRNA-seq data of 10,010 day 7 F-BMDCs (A). Shown are the numbered clusters identified by seurat and subset identities based on expression of key marker genes. UMAP plots on right show expression of the subset annotation markers (gray = low expression; blue = high expression) (right panel). C) Heatmap displaying annotation of cells in the F-BMDCs scRNA-seq dataset using reference gene expression data from the FS tuDC scRNA-seq dataset. Scores represent the proportion of cells within a cluster assigned to a specific cell-type using *SingleR*.

**Figure S4:**
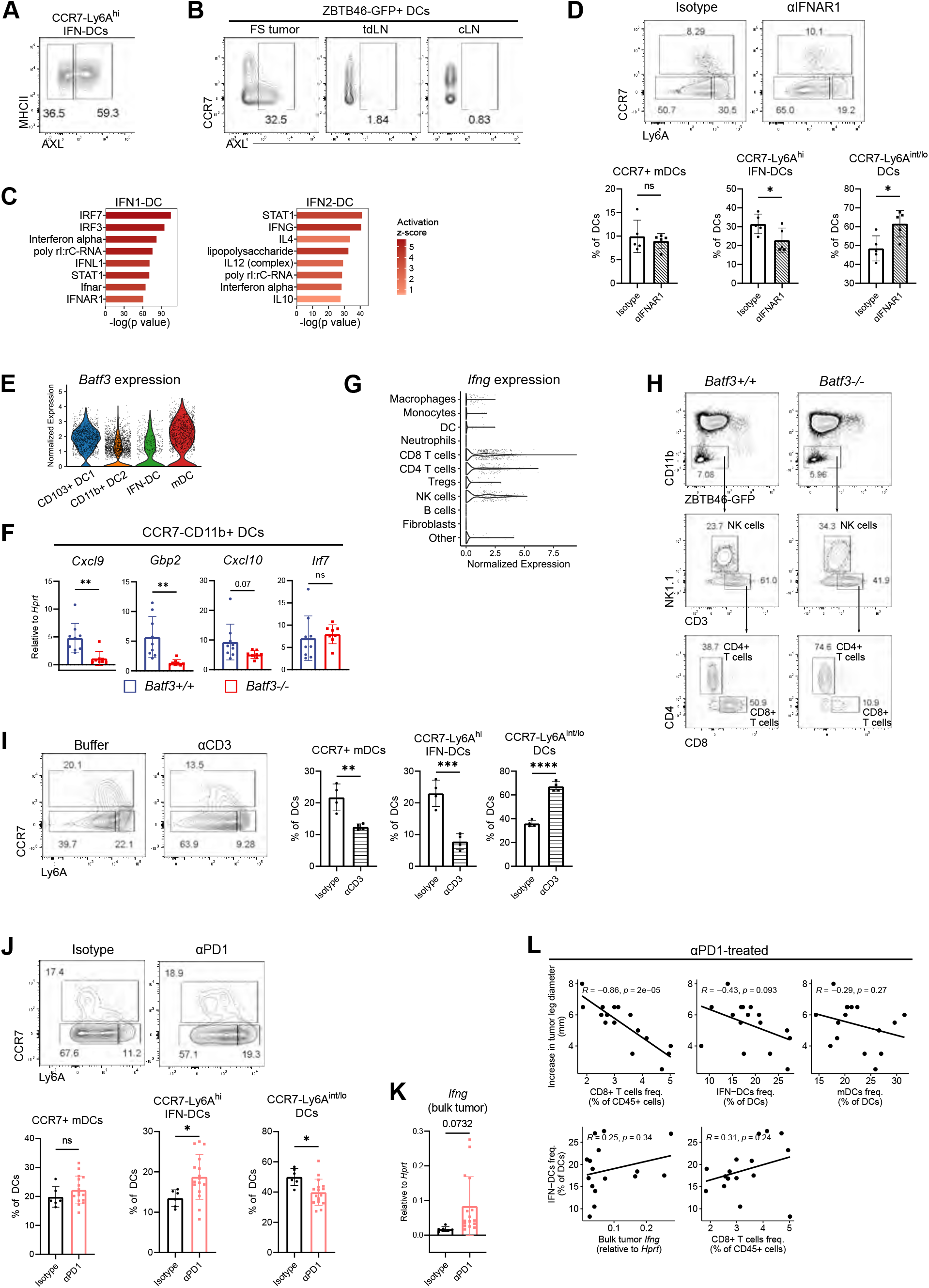
Interferon gamma generates IFN2-DCs. Related to Figure 4. A) FC profiling of AXL staining in CCR7-Ly6A^hi^ IFN-DCs in FS tumors. B) FC profiling of AXL staining in ZBTB46-GFP+ DCs in matched FS tumor, tumor-draining lymph node (tdLN) and non-tumor draining contralateral lymph node (cLN). C) Ingenuity pathway analyses (IPA) predicted upstream regulators of genes differentially expressed in IFN1-DC and IFN2-DC. D) FC profiling of tuDCs from FS tumors treated with control isotype or αIFNAR1 blocking antibody (200 ug intraperitoneal on days 4, 7, 10 and 13 post-tumor injection). Representative FC plots pre-gated on live CD45+CD11C+MHCII+F4/80^lo^Ly6C-tuDCs (left panel). Frequency of tuDC subsets within total tuDCs (right panel). E) Violin plots showing the normalized expression of *Batf3* across the four major tuDC in the FS tuDC scRNA-seq. F) Expression levels (RT-qPCR) of interferon-stimulated genes (ISGs) in sorted CD45+ZBTB46-GFP+ CCR7-CD11B+ tuDCs from FS tumors in *Batf3^+/+^* and *Batf3^-/-^* hosts. G) Violin plots showing the normalized expression of *Ifng* across cell types in a scRNA-seq dataset of CD45+ immune cells from a FS tumor (Devalajara *et al.* 2020). Cell types were annotated based on marker gene expression or with reference to ImmGen expression data via. *SingleR*. H) Flow cytometry profiling of the frequency of lymphoid populations from FS tumors in *Batf3^+/+^* and *Batf3^-/-^* hosts. Events are pre-gated on live CD45+ cells. I) Flow cytometry profiling of tuDCs from FS tumors treated with vehicle or αCD3-depleting antibody (200 ug intraperitoneal on days -3, 0, 3, 6 and 9 post-tumor injection). FS tumors were harvested on day 11. Representative FC plots pre-gated on live CD45+CD11C+MHCII+F4/80^lo^Ly6C-tuDCs (left panel). Frequency of tuDC subsets within total tuDCs (right panel). J-L) FS tumors were treated with control isotype or αPD1 blocking antibody (200 ug intraperitoneal on days 6, 9 and 12 post-tumor injection). J) Flow cytometry profiling of tuDCs. Representative FC plots pre-gated on live CD45+ZBTB46-GFP+ tuDCs (upper panel). Frequency of tuDC subsets within total tuDCs (lower panel). K) Expression levels of bulk tumor *Ifng* as measured by RT-qPCR. L) Correlation within αPD1-treated tumors between tumor growth and the frequency of intratumoral CD8+ T cells (top left), IFN-DCs (top center) or mDCs (top right), or between the frequency of IFN-DCs and the levels of bulk tumor *Ifng* (bottom left) or the frequency of intratumoral CD8+ T cells (bottome right). Correlation coefficients and p values were calculated as the Pearson’s correlation coefficient. \**p* < 0.05, \*\**p* < 0.01, \*\*\**p* < 0.001, \*\*\*\**p* < 0.0001. Statistical analysis was performed with unpaired *t* test (B, G, H, I, and J). Data are shown as mean ± SD.

**Figure S5:**
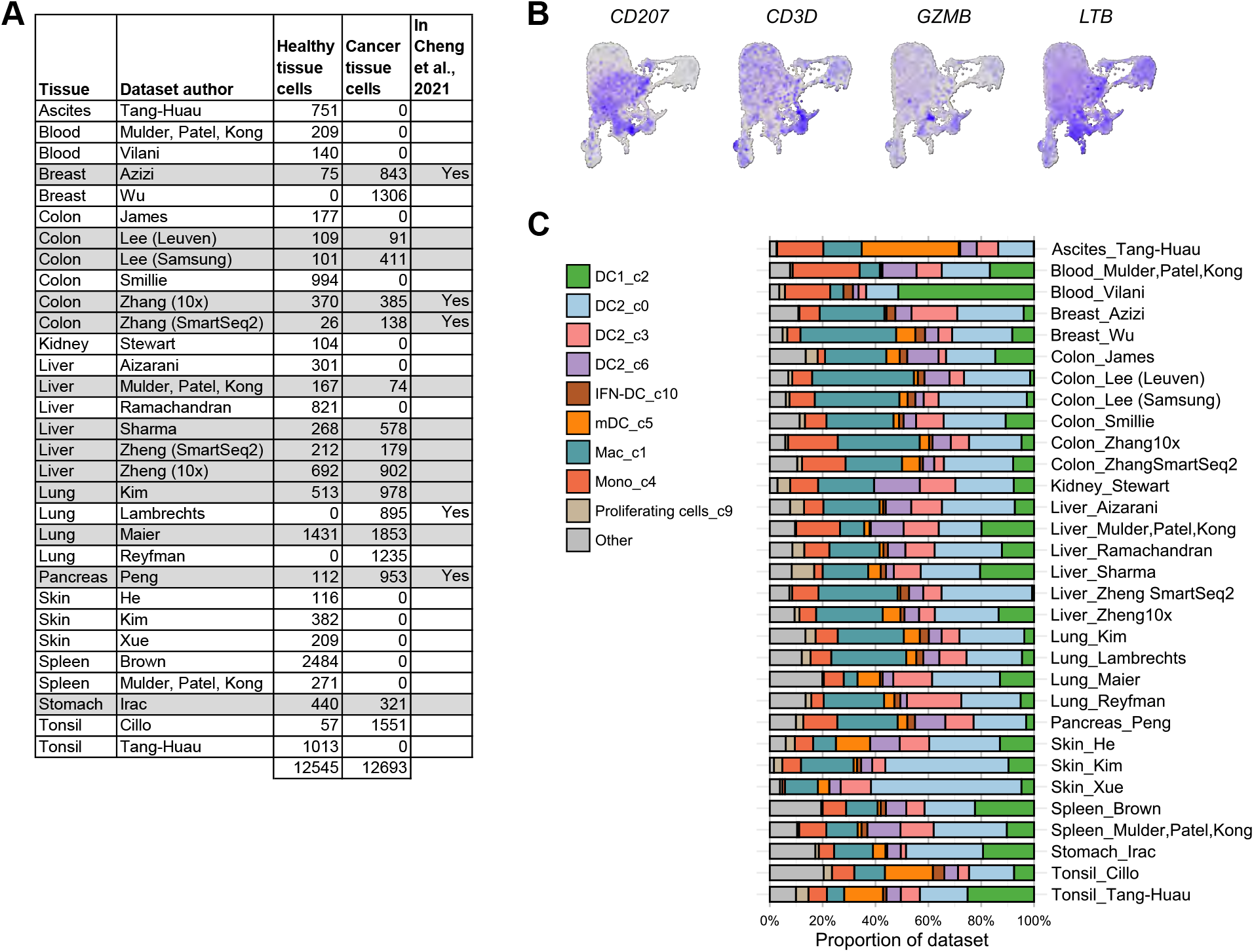
Human DC subsets vary between tissue and tumor.. Related to Figure 5. A) Summary of the datasets used for re-analysis of pre-annotated DCs from the MNP-verse. Datasets in shaded rows were used for comparison of DCs between healthy or cancer status (see Figure 4C and S4D). B) UMAP plots showing expression of genes not associated with cDC subset identity (gray = low expression; blue = high expression). C) Distribution of annotated subsets for each 31 dataset included in the re-analysis of pre-annotated DCs.

## Acknowledgments

We thank the Center for Applied Genomics (CAG) at the Children’s Hospital of Philadelphia (CHOP) for their help and advice on scRNA-seq.

## Funding

This work was supported by the funding sources listed below.

National Institutes of Health grant R37CA234027 (MH)

National Institutes of Health grant F30CA265069 (GPL)

National Institutes of Health grant F30CA236464 (SD)

Burroughs Wellcome grant 1013413.01 (MH)

Concern Foundation (MH)

Cancer Research Institute, Llyod J Old STAR award (MH)

American College of Surgeon Fellowship (IWF)

## Author contributions

Conceptualization: TKJT, MH

Methodology: TKJT, SD, IWF, LZ, GPL, MH

Investigation: TKJT, SD, IWF, MH

Funding acquisition: SD, IWF, GPL, MH

Project administration: MH

Supervision: MH

Writing – original draft: TKJT, MH

Writing – review & editing: TKJT, SD, IWF, LZ, GPL, MH

## Competing interests

The authors declare that they have no competing interests.

## Data and materials availability

**T**he datasets generated in this study are being submitted to the Gene Expression Omnibus (GEO) and the accession numbers will be made available for download.

## Notes

### Competing Interest Statement

The authors have declared no competing interest.

## References

1. A. Mildner, S. Jung, Development and function of dendritic cell subsets. Immunity. 40, 642–656 (2014).

2. M. Guilliams, F. Ginhoux, C. Jakubzick, S. H. Naik, N. Onai, B. U. Schraml, E. Segura, R. Tussiwand, S. Yona, Dendritic cells, monocytes and macrophages: a unified nomenclature based on ontogeny. Nat. Rev. Immunol. 14, 571–578 (2014).

3. M. Merad, P. Sathe, J. Helft, J. Miller, A. Mortha, The dendritic cell lineage: ontogeny and function of dendritic cells and their subsets in the steady state and the inflamed setting. Annu. Rev. Immunol. 31, 563–604 (2013).

4. S. C. Eisenbarth, Dendritic cell subsets in T cell programming: location dictates function. Nat. Rev. Immunol. 19, 89–103 (2019).

5. C. Cheong, I. Matos, J.-H. Choi, D. B. Dandamudi, E. Shrestha, M. P. Longhi, K. L. Jeffrey, R. M. Anthony, C. Kluger, G. Nchinda, H. Koh, A. Rodriguez, J. Idoyaga, M. Pack, K. Velinzon, C. G. Park, R. M. Steinman, Microbial stimulation fully differentiates monocytes to DC-SIGN/CD209(+) dendritic cells for immune T cell areas. Cell. 143, 416– 429 (2010).

6. S. Devalaraja, T. K. J. To, I. W. Folkert, R. Natesan, M. Z. Alam, M. Li, Y. Tada, K. Budagyan, M. T. Dang, L. Zhai, G. P. Lobel, G. E. Ciotti, T. S. K. Eisinger-Mathason, I. A. Asangani, K. Weber, M. C. Simon, M. Haldar, Tumor-Derived Retinoic Acid Regulates Intratumoral Monocyte Differentiation to Promote Immune Suppression. Cell. 180, 1098–1114.e16 (2020).

7. S. K. Wculek, F. J. Cueto, A. M. Mujal, I. Melero, M. F. Krummel, D. Sancho, Dendritic cells in cancer immunology and immunotherapy. Nat. Rev. Immunol. 20, 7–24 (2020).

8. M. L. Broz, M. Binnewies, B. Boldajipour, A. E. Nelson, J. L. Pollack, D. J. Erle, A. Barczak, M. D. Rosenblum, A. Daud, D. L. Barber, S. Amigorena, L. J. Van’t Veer, A. I. Sperling, D. M. Wolf, M. F. Krummel, Dissecting the tumor myeloid compartment reveals rare activating antigen-presenting cells critical for T cell immunity. Cancer Cell. 26, 638– 652 (2014).

9. K. Hildner, B. T. Edelson, W. E. Purtha, M. Diamond, H. Matsushita, M. Kohyama, B. Calderon, B. U. Schraml, E. R. Unanue, M. S. Diamond, R. D. Schreiber, T. L. Murphy, K. M. Murphy, Batf3 deficiency reveals a critical role for CD8alpha+ dendritic cells in cytotoxic T cell immunity. Science. 322, 1097–1100 (2008).

10. E. Papalexi, R. Satija, Single-cell RNA sequencing to explore immune cell heterogeneity. Nat. Rev. Immunol. 18, 35–45 (2018).

11. G. M. Gerhard, R. Bill, M. Messemaker, A. M. Klein, M. J. Pittet, Tumor-infiltrating dendritic cell states are conserved across solid human cancers. J Exp Med. 218 (2021), doi:10.1084/jem.20200264.

12. A. Chow, B. D. Brown, M. Merad, Studying the mononuclear phagocyte system in the molecular age. Nat Rev Immunol. 11, 788–798 (2011).

13. C. L. Abram, G. L. Roberge, Y. Hu, C. A. Lowell, Comparative analysis of the efficiency and specificity of myeloid-Cre deleting strains using ROSA-EYFP reporter mice. J Immunol Methods. 408, 89–100 (2014).

14. A. T. Satpathy, W. Kc, J. C. Albring, B. T. Edelson, N. M. Kretzer, D. Bhattacharya, T. L. Murphy, K. M. Murphy, Zbtb46 expression distinguishes classical dendritic cells and their committed progenitors from other immune lineages. J Exp Med. 209, 1135–1152 (2012).

15. D. Laoui, E. Van Overmeire, K. Movahedi, J. Van den Bossche, E. Schouppe, C. Mommer, A. Nikolaou, Y. Morias, P. De Baetselier, J. A. Van Ginderachter, Mononuclear phagocyte heterogeneity in cancer: different subsets and activation states reaching out at the tumor site. Immunobiology. 216, 1192–1202 (2011).

16. J. Sheng, Q. Chen, I. Soncin, S. L. Ng, K. Karjalainen, C. Ruedl, A Discrete Subset of Monocyte-Derived Cells among Typical Conventional Type 2 Dendritic Cells Can Efficiently Cross-Present. Cell Rep. 21, 1203–1214 (2017).

17. Y. Hao, S. Hao, E. Andersen-Nissen, W. M. Mauck, S. Zheng, A. Butler, M. J. Lee, A. J. Wilk, C. Darby, M. Zager, P. Hoffman, M. Stoeckius, E. Papalexi, E. P. Mimitou, J. Jain, A. Srivastava, T. Stuart, L. M. Fleming, B. Yeung, A. J. Rogers, J. M. McElrath, C. A. Blish, R. Gottardo, P. Smibert, R. Satija, Integrated analysis of multimodal single-cell data. Cell. 184, 3573–3587.e29 (2021).

18. J. C. Miller, B. D. Brown, T. Shay, E. L. Gautier, V. Jojic, A. Cohain, G. Pandey, M. Leboeuf, K. G. Elpek, J. Helft, D. Hashimoto, A. Chow, J. Price, M. Greter, M. Bogunovic, A. Bellemare-Pelletier, P. S. Frenette, G. J. Randolph, S. J. Turley, M. Merad, Immunological Genome Consortium, Deciphering the transcriptional network of the dendritic cell lineage. Nat Immunol. 13, 888–899 (2012).

19. B. Maier, A. M. Leader, S. T. Chen, N. Tung, C. Chang, J. LeBerichel, A. Chudnovskiy, S. Maskey, L. Walker, J. P. Finnigan, M. E. Kirkling, B. Reizis, S. Ghosh, N. R. D’Amore, N. Bhardwaj, C. V. Rothlin, A. Wolf, R. Flores, T. Marron, A. H. Rahman, E. Kenigsberg, B. D. Brown, M. Merad, A conserved dendritic-cell regulatory program limits antitumour immunity. Nature. 580, 257–262 (2020).

20. Q. Zhang, Y. He, N. Luo, S. J. Patel, Y. Han, R. Gao, M. Modak, S. Carotta, C. Haslinger, D. Kind, G. W. Peet, G. Zhong, S. Lu, W. Zhu, Y. Mao, M. Xiao, M. Bergmann, X. Hu, S. P. Kerkar, A. B. Vogt, S. Pflanz, K. Liu, J. Peng, X. Ren, Z. Zhang, Landscape and Dynamics of Single Immune Cells in Hepatocellular Carcinoma. Cell. 179, 829–845.e20 (2019).

21. R. Zilionis, C. Engblom, C. Pfirschke, V. Savova, D. Zemmour, H. D. Saatcioglu, I. Krishnan, G. Maroni, C. V. Meyerovitz, C. M. Kerwin, S. Choi, W. G. Richards, A. De Rienzo, D. G. Tenen, R. Bueno, E. Levantini, M. J. Pittet, A. M. Klein, Single-Cell Transcriptomics of Human and Mouse Lung Cancers Reveals Conserved Myeloid Populations across Individuals and Species. Immunity. 50, 1317–1334.e10 (2019).

22. L. Ardouin, H. Luche, R. Chelbi, S. Carpentier, A. Shawket, F. Montanana Sanchis, C. Santa Maria, P. Grenot, Y. Alexandre, C. Grégoire, A. Fries, T.-P. Vu Manh, S. Tamoutounour, K. Crozat, E. Tomasello, A. Jorquera, E. Fossum, B. Bogen, H. Azukizawa, M. Bajenoff, S. Henri, M. Dalod, B. Malissen, Broad and Largely Concordant Molecular Changes Characterize Tolerogenic and Immunogenic Dendritic Cell Maturation in Thymus and Periphery. Immunity. 45, 305–318 (2016).

23. M. Dalod, R. Chelbi, B. Malissen, T. Lawrence, Dendritic cell maturation: functional specialization through signaling specificity and transcriptional programming. EMBO J. 33, 1104–1116 (2014).

24. M. E. Mikucki, D. T. Fisher, J. Matsuzaki, J. J. Skitzki, N. B. Gaulin, J. B. Muhitch, A. W. Ku, J. G. Frelinger, K. Odunsi, T. F. Gajewski, A. D. Luster, S. S. Evans, Non-redundant requirement for CXCR3 signalling during tumoricidal T-cell trafficking across tumour vascular checkpoints. Nat Commun. 6, 7458 (2015).

25. D. Dangaj, M. Bruand, A. J. Grimm, C. Ronet, D. Barras, P. A. Duttagupta, E. Lanitis, J. Duraiswamy, J. L. Tanyi, F. Benencia, J. Conejo-Garcia, H. R. Ramay, K. T. Montone, D. J. Powell, P. A. Gimotty, A. Facciabene, D. G. Jackson, J. S. Weber, S. J. Rodig, S. F. Hodi, L. E. Kandalaft, M. Irving, L. Zhang, P. Foukas, S. Rusakiewicz, M. Delorenzi, G. Coukos, Cooperation between Constitutive and Inducible Chemokines Enables T Cell Engraftment and Immune Attack in Solid Tumors. Cancer Cell. 35, 885–900.e10 (2019).

26. M. Guilliams, C.-A. Dutertre, C. L. Scott, N. McGovern, D. Sichien, S. Chakarov, S. Van Gassen, J. Chen, M. Poidinger, S. De Prijck, S. J. Tavernier, I. Low, S. E. Irac, C. N. Mattar, H. R. Sumatoh, G. H. L. Low, T. J. K. Chung, D. K. H. Chan, K. K. Tan, T. L. K. Hon, E. Fossum, B. Bogen, M. Choolani, J. K. Y. Chan, A. Larbi, H. Luche, S. Henri, Y. Saeys, E. W. Newell, B. N. Lambrecht, B. Malissen, F. Ginhoux, Unsupervised High-Dimensional Analysis Aligns Dendritic Cells across Tissues and Species. Immunity. 45, 669–684 (2016).

27. P. Michea, F. Noël, E. Zakine, U. Czerwinska, P. Sirven, O. Abouzid, C. Goudot, A. Scholer-Dahirel, A. Vincent-Salomon, F. Reyal, S. Amigorena, M. Guillot-Delost, E. Segura, V. Soumelis, Adjustment of dendritic cells to the breast-cancer microenvironment is subset specific. Nat Immunol. 19, 885–897 (2018).

28. S. H. Naik, A. I. Proietto, N. S. Wilson, A. Dakic, P. Schnorrer, M. Fuchsberger, M. H. Lahoud, M. O’Keeffe, Q. Shao, W. Chen, J. A. Villadangos, K. Shortman, L. Wu, Cutting edge: generation of splenic CD8+ and CD8-dendritic cell equivalents in Fms-like tyrosine kinase 3 ligand bone marrow cultures. J Immunol. 174, 6592–6597 (2005).

29. C. Bosteels, K. Neyt, M. Vanheerswynghels, M. J. van Helden, D. Sichien, N. Debeuf, S. De Prijck, V. Bosteels, N. Vandamme, L. Martens, Y. Saeys, E. Louagie, M. Lesage, D. L. Williams, S.-C. Tang, J. U. Mayer, F. Ronchese, C. L. Scott, H. Hammad, M. Guilliams, B. N. Lambrecht, Inflammatory Type 2 cDCs Acquire Features of cDC1s and Macrophages to Orchestrate Immunity to Respiratory Virus Infection. Immunity. 52, 1039–1056.e9 (2020).

30. G. Ghislat, A. S. Cheema, E. Baudoin, C. Verthuy, P. J. Ballester, K. Crozat, N. Attaf, C. Dong, P. Milpied, B. Malissen, N. Auphan-Anezin, T. P. V. Manh, M. Dalod, T. Lawrence, NF-κB-dependent IRF1 activation programs cDC1 dendritic cells to drive antitumor immunity. Sci Immunol. 6, eabg3570 (2021).

31. E. Duong, T. B. Fessenden, E. Lutz, T. Dinter, L. Yim, S. Blatt, A. Bhutkar, K. D. Wittrup, S. Spranger, Type I interferon activates MHC class I-dressed CD11b+ conventional dendritic cells to promote protective anti-tumor CD8+ T cell immunity. Immunity, S1074–7613(21)00458–1 (2021).

32. P. C. Tumeh, C. L. Harview, J. H. Yearley, I. P. Shintaku, E. J. M. Taylor, L. Robert, B. Chmielowski, M. Spasic, G. Henry, V. Ciobanu, A. N. West, M. Carmona, C. Kivork, E. Seja, G. Cherry, A. J. Gutierrez, T. R. Grogan, C. Mateus, G. Tomasic, J. A. Glaspy, R. O. Emerson, H. Robins, R. H. Pierce, D. A. Elashoff, C. Robert, A. Ribas, PD-1 blockade induces responses by inhibiting adaptive immune resistance. Nature. 515, 568–571 (2014).

33. C.-A. Dutertre, L.-F. Wang, F. Ginhoux, Aligning bona fide dendritic cell populations across species. Cell Immunol. 291, 3–10 (2014).

34. K. Mulder, A. A. Patel, W. T. Kong, C. Piot, E. Halitzki, G. Dunsmore, S. Khalilnezhad, S. E. Irac, A. Dubuisson, M. Chevrier, X. M. Zhang, J. K. C. Tam, T. K. H. Lim, R. M. M. Wong, R. Pai, A. I. S. Khalil, P. K. H. Chow, S. Z. Wu, G. Al-Eryani, D. Roden, A. Swarbrick, J. K. Y. Chan, S. Albani, L. Derosa, L. Zitvogel, A. Sharma, J. Chen, A. Silvin, A. Bertoletti, C. Blériot, C.-A. Dutertre, F. Ginhoux, Cross-tissue single-cell landscape of human monocytes and macrophages in health and disease. Immunity. 54, 1883–1900.e5 (2021).

35. S. Cheng, Z. Li, R. Gao, B. Xing, Y. Gao, Y. Yang, S. Qin, L. Zhang, H. Ouyang, P. Du, L. Jiang, B. Zhang, Y. Yang, X. Wang, X. Ren, J.-X. Bei, X. Hu, Z. Bu, J. Ji, Z. Zhang, A pan-cancer single-cell transcriptional atlas of tumor infiltrating myeloid cells. Cell. 184, 792–809.e23 (2021).

36. S. Tamoutounour, M. Guilliams, F. Montanana Sanchis, H. Liu, D. Terhorst, C. Malosse, E. Pollet, L. Ardouin, H. Luche, C. Sanchez, M. Dalod, B. Malissen, S. Henri, Origins and functional specialization of macrophages and of conventional and monocyte-derived dendritic cells in mouse skin. Immunity. 39, 925–938 (2013).

37. E. W. Roberts, M. L. Broz, M. Binnewies, M. B. Headley, A. E. Nelson, D. M. Wolf, T. Kaisho, D. Bogunovic, N. Bhardwaj, M. F. Krummel, Critical Role for CD103(+)/CD141(+) Dendritic Cells Bearing CCR7 for Tumor Antigen Trafficking and Priming of T Cell Immunity in Melanoma. Cancer Cell. 30, 324–336 (2016).

38. S. Spranger, D. Dai, B. Horton, T. F. Gajewski, Tumor-Residing Batf3 Dendritic Cells Are Required for Effector T Cell Trafficking and Adoptive T Cell Therapy. Cancer Cell. 31, 711–723.e4 (2017).

39. Á. de Mingo Pulido, A. Gardner, S. Hiebler, H. Soliman, H. S. Rugo, M. F. Krummel, L. M. Coussens, B. Ruffell, TIM-3 Regulates CD103+ Dendritic Cell Function and Response to Chemotherapy in Breast Cancer. Cancer Cell. 33, 60–74.e6 (2018).

40. M. T. Chow, A. J. Ozga, R. L. Servis, D. T. Frederick, J. A. Lo, D. E. Fisher, G. J. Freeman, G. M. Boland, A. D. Luster, Intratumoral Activity of the CXCR3 Chemokine System Is Required for the Efficacy of Anti-PD-1 Therapy. Immunity. 50, 1498–1512.e5 (2019).

41. I. G. House, P. Savas, J. Lai, A. X. Y. Chen, A. J. Oliver, Z. L. Teo, K. L. Todd, M. A. Henderson, L. Giuffrida, E. V. Petley, K. Sek, S. Mardiana, T. N. Gide, C. Quek, R. A. Scolyer, G. V. Long, J. S. Wilmott, S. Loi, P. K. Darcy, P. A. Beavis, Macrophage-Derived CXCL9 and CXCL10 Are Required for Antitumor Immune Responses Following Immune Checkpoint Blockade. Clin Cancer Res. 26, 487–504 (2020).

42. C. S. Garris, S. P. Arlauckas, R. H. Kohler, M. P. Trefny, S. Garren, C. Piot, C. Engblom, C. Pfirschke, M. Siwicki, J. Gungabeesoon, G. J. Freeman, S. E. Warren, S. Ong, E. Browning, C. G. Twitty, R. H. Pierce, M. H. Le, A. P. Algazi, A. I. Daud, S. I. Pai, A. Zippelius, R. Weissleder, M. J. Pittet, Successful Anti-PD-1 Cancer Immunotherapy Requires T Cell-Dendritic Cell Crosstalk Involving the Cytokines IFN-γ and IL-12. Immunity. 49, 1148–1161.e7 (2018).

43. M. Di Pilato, R. Kfuri-Rubens, J. N. Pruessmann, A. J. Ozga, M. Messemaker, B. L. Cadilha, R. Sivakumar, C. Cianciaruso, R. D. Warner, F. Marangoni, E. Carrizosa, S. Lesch, J. Billingsley, D. Perez-Ramos, F. Zavala, E. Rheinbay, A. D. Luster, M. Y. Gerner, S. Kobold, M. J. Pittet, T. R. Mempel, CXCR6 positions cytotoxic T cells to receive critical survival signals in the tumor microenvironment. Cell. 184, 4512–4530.e22 (2021).

